# Flaviviruses alter endoplasmic reticulum-mitochondria contacts to regulate respiration and apoptosis

**DOI:** 10.1101/2023.03.09.531853

**Authors:** Wesley Freppel, Anaïs Anton, Zaynab Nouhi, Clément Mazeaud, Claudia Gilbert, Nicolas Tremblay, Viviana Andrea Barragan Torres, Aïssatou Aïcha Sow, Xavier Laulhé, Alain Lamarre, Ian Gaël Rodrigue-Gervais, Andreas Pichlmair, Pietro Scaturro, Laura Hulea, Laurent Chatel-Chaix

**Author notes:** Current address: Institute for Glycomics, Gold Coast Campus, Griffith University, Southport, QLD 4222, Australia. Corresponding author: Laurent Chatel-Chaix.

## Abstract

With no therapeutics available, there is an urgent need to better understand the pathogenesis of flaviviruses which constitute a threat to public health worldwide. During infection, dengue virus (DENV) and Zika virus (ZIKV), two flaviviruses induce alterations of mitochondria morphology to favor viral replication, suggesting a viral co-opting of mitochondria functions. Here, we performed an extensive transmission electron microscopy-based quantitative analysis to demonstrate that both DENV and ZIKV alter endoplasmic reticulum-mitochondria contacts (ERMC). This correlated at the molecular level with an impairment of ERMC tethering protein complexes located at the surface of both organelles. Furthermore, virus infection, as well as NS4B expression modulated the mitochondrial oxygen consumption rate. Consistently, metabolomic and mitoproteomic analyses revealed a decrease in the abundance of several metabolites of the Krebs cycle and changes in the stoichiometry of the electron transport chain. Most importantly, ERMC destabilization by protein knockdown increased virus replication while dampening ZIKV-induced apoptosis. Overall, our results support the notion that flaviviruses hijack ERMCs to generate a cytoplasmic environment beneficial for sustained and efficient replication.

## INTRODUCTION

Flavivirus infections constitute a major public health concern worldwide. With an estimation of 390 million people infected per year, dengue virus (DENV) causes the most prevalent arthropod-borne viral disease (Bhatt *et al*, 2013). Although DENV mainly circulates in (sub-)tropical regions, it has also now reached Europe and North America because of vector colonization of these areas (Liu-Helmersson *et al*, 2016; Ryan *et al*, 2019). Upon infection through the bite of an *Aedes*-type mosquito, symptoms associated with dengue fever can manifest as severe fevers which may be haemorrhagic and eventually lead to shock syndrome and death (WHO, 2022). In 2015, Zika virus (ZIKV), another flavivirus closely related to DENV, has quickly emerged in South America causing unexpected symptoms such as microcephaly in newborns following congenital transmission, in addition to other neurological complications in adults such as Guillain-Barré syndrome (Carod-Artal, 2018; Moore *et al*, 2017). Unfortunately, no treatments against DENV and ZIKV are currently available, partly due to our limited knowledge of the cellular and molecular mechanisms involved in flavivirus life cycle and pathogenesis, which could constitute therapeutic targets.

DENV and ZIKV are single positive-stranded RNA viruses belonging to the *Flavivirus* genus within the *Flaviviridae* family. Following entry into the host cell, the viral RNA genome (vRNA) is translated into a single polyprotein at the membrane of the endoplasmic reticulum (ER). This viral protein product is cleaved by host and viral proteases, generating three structural proteins (C, prM and E) which assemble with the vRNA to form new viral particles, and into 7 non-structural proteins (NS1, NS2A, NS2B, NS3, NS4A, NS4B, NS5) which are involved in the replication of the viral genome (Mazeaud *et al*, 2018). The viral replication takes place in cytoplasmic substructures called viral replication organelles (vRO) that are derived from ER membrane alterations induced by the virus (Chatel-Chaix & Bartenschlager, 2014; Cortese *et al*, 2017; Gillespie *et al*, 2010; Junjhon *et al*, 2014; Miorin *et al*, 2013; Paul & Bartenschlager, 2015; Welsch *et al*, 2009). vROs comprise: 1-vesicle packets, believed to be the site of vRNA replication; 2-virus bags, ER cisternae in which immature assembled virions accumulate; and 3-convoluted membranes (CM), that are enriched in NS3, NS4B, and NS4A (Chatel-Chaix *et al*, 2016; Miller *et al*, 2007; Welsch *et al*., 2009) whose functions are poorly understood. It was proposed that CMs dampen antiviral cellular processes, such as early innate immunity response and apoptosis to favor viral replication.

In DENV-and ZIKV-infected cells, mitochondria make physical contacts with CMs and exhibit an elongated morphology, which stimulates viral replication (Barbier *et al*, 2017; Chatel-Chaix *et al*., 2016). This regulation of mitochondrial morphodynamics was attributed to NS4B which partly resides in CMs. The pharmacological destabilization of CMs induces mitochondria fragmentation and correlates with a stimulation of virus-induced apoptosis (Anton *et al*, 2021). Conversely, mitochondria elongation positively influences the size and abundance of CMs and dampens the RIG-I-dependent type-I and −III interferon induction (Chatel-Chaix *et al*., 2016). This supports a model in which flaviviruses regulate mitochondrial functions through their contacts with CMs for the benefit of replication.

ER-mitochondria contacts (ERMC) rely on protein-protein connections maintaining a 10-25 nm-wide interface between the ER and the mitochondria that allows molecular transfers between both organelles (Csordas *et al*, 2006; Fujimoto & Hayashi, 2011; Vance, 2015). This ultrastructure contributes to several cellular processes such as calcium homeostasis, lipid transport, autophagy regulation, mitochondrial morphodynamics, apoptosis induction and early innate immunity (Cohen *et al*, 2018; Friedman *et al*, 2011; Hamasaki *et al*, 2013; Pourcelot & Arnoult, 2014; Schwarz & Blower, 2016). Illustrating their important roles in cellular processes, ERMC are a target of viruses for interfering with mitochondrial-mediated antiviral responses (Horner *et al*, 2011; Horner *et al*, 2015). Recently, we have shown that DENV-induced mitochondria elongation positively regulates the biogenesis of CM (Chatel-Chaix *et al*., 2016).. This is accompanied with a decrease in colocalization between the ER and mitochondria at the cellular level in confocal microscopy. However, it is still unclear whether flaviviruses globally alter ERMC and what the resulting impacts are regarding cellular processes such as oxidative respiration and apoptosis.

In this study, we show an overall alteration of the ERMC compartment following DENV and ZIKV infections. Concomitantly, reducing expression of several ERMC proteins responsible for tethering mitochondria and the ER increased DENV and ZIKV replication, supporting that both viruses alter ERMC to stimulate viral replication. Interestingly, the expression profile of several ERMC proteins was changed with a drastic decrease in the levels of RRBP1 over the course of the infection and the appearance of an alternative ZIKV-specific SYNJ2BP protein product. This suggests that both viruses destabilize ERMC by targeting tethering proteins. Furthermore, both DENV and ZIKV modulated mitochondrial respiratory metabolism in living cells. This correlated with a decrease in the abundance of several metabolites of the Krebs cycle and changes in the stoichiometry of the electron transport chain. Most importantly, targeting ERMC by silencing either RRBP1 or SYNJ2BP increased respiration and dampened ZIKV-induced apoptosis supporting the importance of ERMC alteration by DENV and ZIKV for attenuating antiviral cellular processes and for maintaining a cytoplasmic environment favorable to the viral replication.

## RESULTS

### DENV and ZIKV alter ERMCs

Previous observation of DENV-infected Huh7 cells at the ultrastructural level suggested that ERMCs were altered. Although this was not quantified, this correlated with a decrease of 3D colocalization between mitochondria and ER in a limited number of analyzed cells in confocal microscopy, *i.e.*, at a resolution which is too low to precisely measure heterotypic organelle contacts (Chatel-Chaix *et al*., 2016). The molecular mechanisms underlying this phenotype was not described and it was also not established whether this phenotype is specific for DENV or also expands to other relevant flaviviruses, such as ZIKV. To clearly address whether DENV and ZIKV infections induce a global alteration of ERMCs, we performed a comprehensive quantitative ultrastructural analysis of DENV-and ZIKV-infected Huh7.5 hepatoma cells using transmission electron microscopy. The Huh7.5 cell line was chosen as a model because these cells are defective in RIG-I-dependent interferon induction, warranting that any observed phenotype is not caused by early innate immunity, a process that relies on mitochondrial protein MAVS (Loo & Gale, 2011). More than two hundred mitochondria per conditions (n=215-1. 351) were analyzed for ERMCs following 48 hrs of infection with the ZIKV H/PF/2013 contemporary strain or serotype 2 DENV 16681s strain. In uninfected conditions, 74.7% of mitochondria were surrounded by ER tubules with up to 75% of the mitochondrial perimeter being in contact with these ER membranes (median at 31.4%; Fig. 1a-c). In stark contrast, both DENV and ZIKV-infected cells exhibited a reduced proportion of mitochondria in contact with ER membranes, with a phenotype more pronounced for DENV than for ZIKV. In addition, when the two organelles were still physically associated, only 19% of the mitochondria perimeter in average were in contact with ER following both ZIKV and DENV infections, compared to 35% in uninfected condition (Fig 1c). Consistent with the previous observation, mitochondria showed an elongated morphology and were often located in the vicinity of CMs (Fig. 1a).

**Figure 1:**
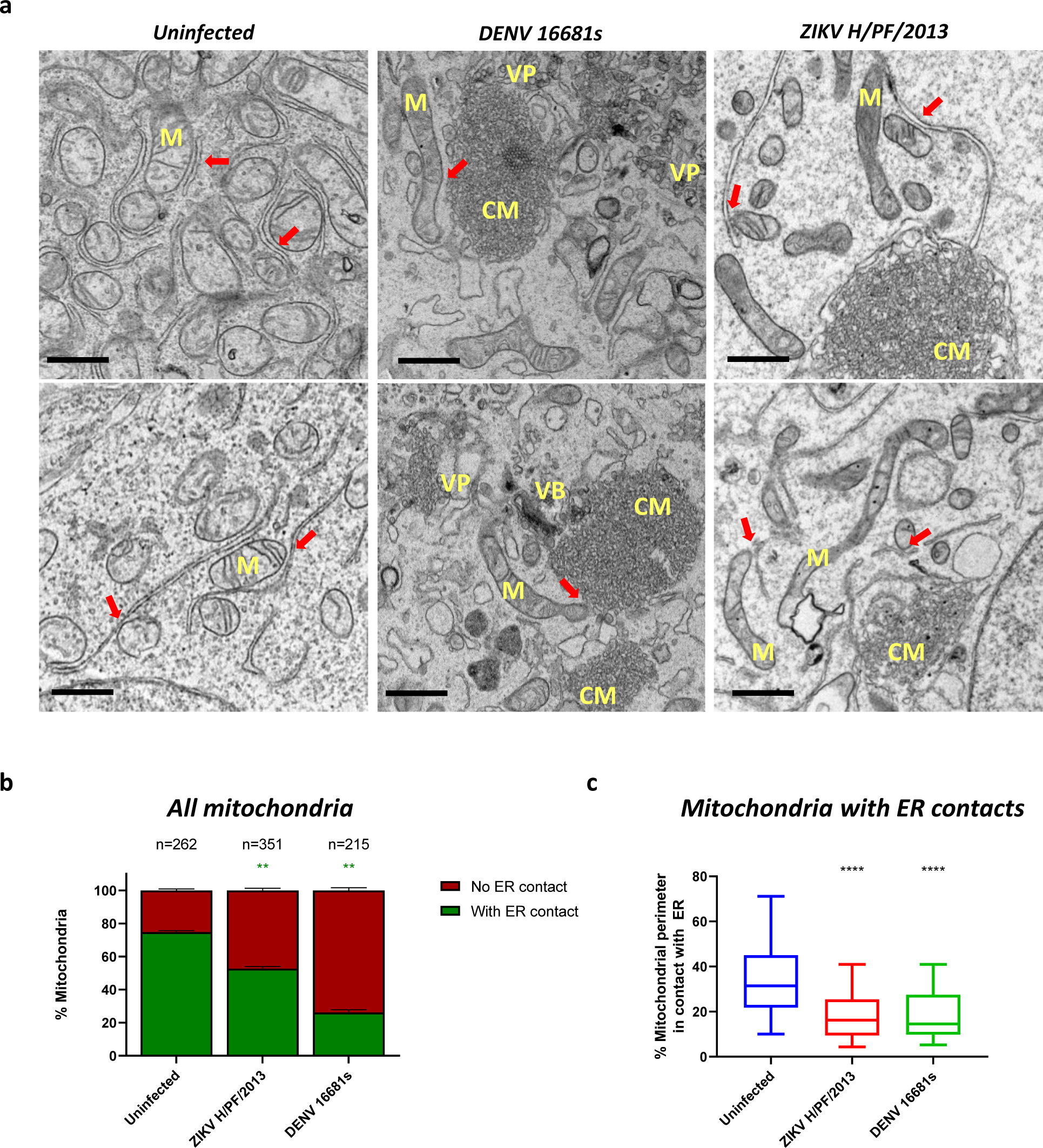
DENV and ZIKV infection alter endoplasmic reticulum-mitochondria contact sites. Huh7.5 cells were infected with DENV2 16681s (multiplicity of infection (MOI)=1), or ZIKV H/PF/2013 (MOI=10) or left uninfected. Forty-eight hours later, cells were processed for transmission electron microscopy. (a) Electron micrographs of uninfected and infected cells. Red arrows indicate ER-mitochondria contact sites. CM: convoluted membranes; VP: vesicle packets; M: mitochondria; VB: virus bags. Scale bar = 1 μm. (b) Proportion of mitochondria with or without contacts with the ER; n: number of analyzed mitochondria. (c) Percentage of the mitochondrial perimeter in contact with ER. The mitochondria of (b) which were not in contact with ER were excluded from this quantification. Values in the 5%-95% percentile range are shown. Analyses were made with micrographs from 2 independent experiments.

To examine the impact of both virus infections on ERMCs at a molecular level in a larger cell population, we analyzed the extent of protein-protein interactions involved in tethering mitochondria and ER using proximity ligation assay (PLA). This technique allows the detection of the proximity between two proteins at a maximum distance of 40 nm and the analysis of the intracellular localization of protein complexes using confocal microscopy. We focused our PLA analysis on three ERMC tethering complexes known to contribute to ERMC formation and/or stability, namely the interactions between: 1-ER −resident vesicle-associated membrane protein-associated protein B (VAPB) and the mitochondrial outer membrane protein, protein tyrosine phosphatase-interacting protein 51 (PTPIP51) (De Vos *et al*, 2012); 2-ER-resident, ribosome-binding protein 1 (RRBP1) and the mitochondrial protein, synaptojanin 2-binding protein (SYNJ2BP) (Duan *et al*, 2022; Hung *et al*, 2019); and 3-the ER protein, inositol 1,4,5-trisphosphate receptor 1 (IP3R1) and mitochondrial voltage-dependent anion-selective channel 1 (VDAC1) via the chaperone glucose-regulated protein 75 (GRP75) (Hartmann & Verkhratsky, 1998) (Fig. 2a). First, we confirmed that PLA signals detected for these three different tethering complexes mostly colocalized with the mitochondria network in Huh7.5 cells stably expressing the mitochondria-localized mito-mTurquoise2 fluorophore (Suppl. Fig. 1a). PLAs were then performed for VAPB-PTPIP51, RRBP1-SYNJ2PB and IP3R1-VDAC1 interactions in uninfected-or DENV/ZIKV infected-Huh7.5 cells at 48 and 72 hrs post-infection and combined with immunostaining of NS3 viral protein to identify infected cells. As expected, we detected a high amount of RRBP1-SYNJ2BP (Fig. 2b), IP3R1-VDAC1 (Suppl. Fig. 1b) and VAPB-PTPIP51 (Suppl. Fig. 1c) interactions in the uninfected condition in confocal microscopy. The signal was specific to the targeted protein-protein interactions since omitting either primary antibody gave very little, if any, PLA signal (Fig. 2c, Ab alone controls). Strikingly, when comparing ZIKV-or DENV-infected (green) and uninfected cells in the same images, it was obvious that the RRBP1-SYNJ2BP PLA signal was less abundant in infected cells (Fig 2b). Subsequent quantification of the PLA dots per cell demonstrated an overall decrease in PLA signal in NS3-positive cells *(i.e.*, infected) in all three assays (Fig. 2c, Suppl. Fig. 1b-c). This demonstrates a decrease in the abundance of the RRBP1-SYNJ2BP, IP3R1-VDAC1 and VABP-PTPIP51 ERMC tethering complexes and confirms an alteration of ERMCs by DENV and ZIKV at the molecular level. Interestingly, this decrease was particularly prominent for RRBP1-SYNJ2BP and IP3R1-VDAC1 at 72 hrs post-infection.

**Figure 2:**
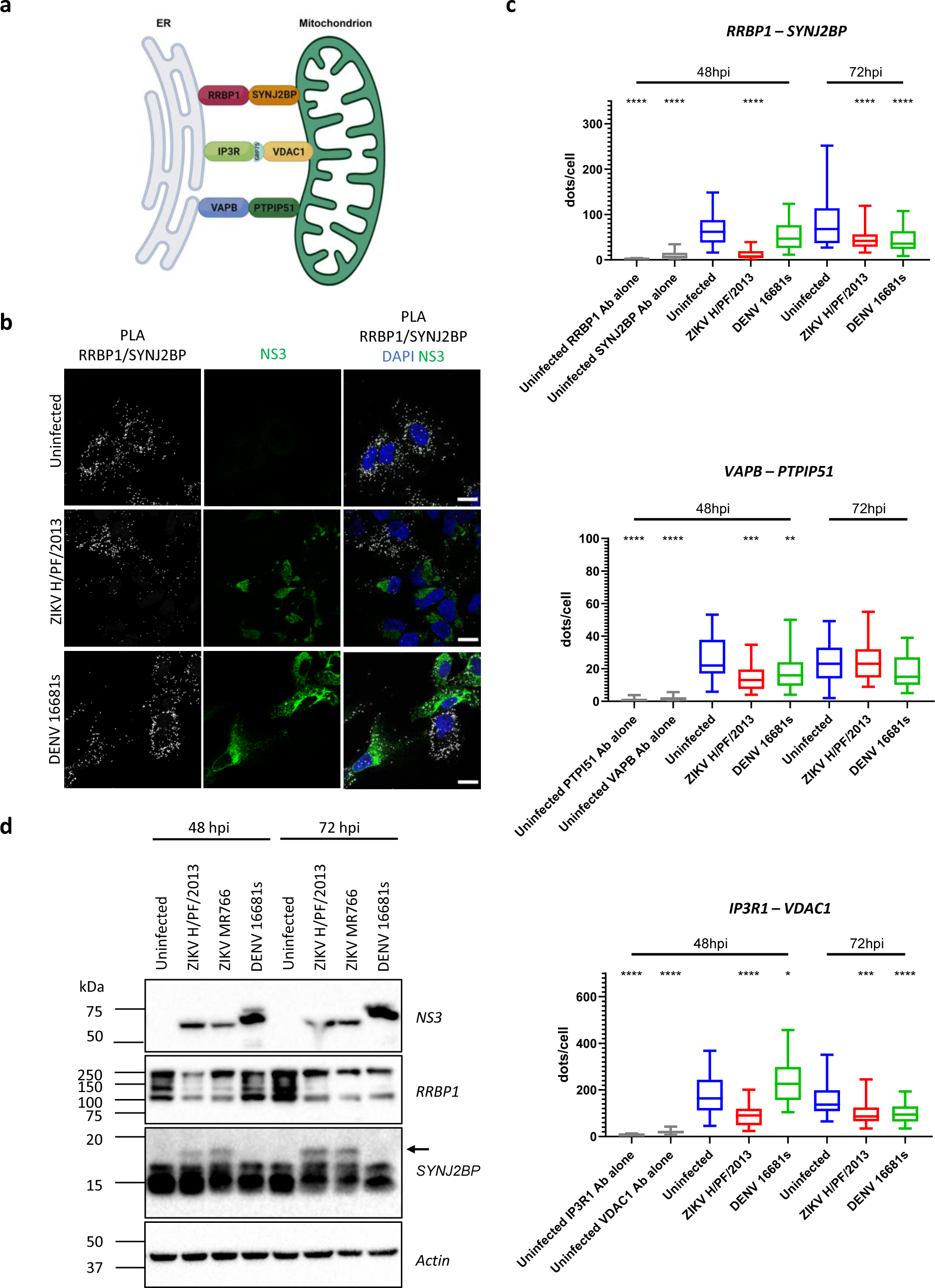
Impact of DENV and ZIKV infection on the expression of ERMC proteins. (a) Schematic representation of ERMC tethering complexes. Generated with BioRender. (b) Huh7.5 cells were infected with DENV 16681s (MOI=1) or ZIKV H/PF/2013 (MOI=10) or left uninfected. Two or three days later, cells were fixed and subjected to proximity ligation assays (PLA) to detect SYNJ2BP-RRBP1 interactions, and immunostained for NS3 viral protein to identify infected cells. Cells were imaged using confocal microscopy. Representative images are shown. Scale bar=20 µm. (c) Quantification of PLA dot abundance for RRBP1-SYNJ2BP, VAPB-PTPIP51 and IP3R1-VDAC1 interactions in uninfected-and infected-cells from two independent experiments. Values in the 5%-95% percentile range are shown. (d) Cells were prepared as in (b). Two or three days post-infection, cell extracts were prepared, and the expression levels of the indicated proteins were analyzed by western blotting. The arrow indicates the ZIKV-induced SYNJ2BP protein product.

To examine the impact of DENV and ZIKV infection on the expression of ERMC proteins, we performed western blotting on extracts from uninfected or infected Huh7.5 cells at 48 and 72hrs post-infection. The expression levels of GRP75, VDAC1, VAPB and PTPIP51 were unchanged upon ZIKV and DENV infections (Suppl. Fig. 2a-b). In contrast, the expression of RRBP1 was reduced at 48 hrs post-infection with a further decrease of its levels at 72hrs post-infection (Fig. 2d and Suppl. Fig. 2a). Moreover, an alternative SYNJ2BP product exhibiting a higher molecular weight was detected in ZIKV-infected cells at 48 and 72 hrs post-infection (Fig. 2d and Suppl. Fig. 2a). Finally, our immunofluorescence-based confocal microscopy analysis of DENV-and ZIKV-infected cells did not reveal any changes in the sub-cellular distribution of IP3R1, VDAC1, RRBP1, SYNJ2BP and PTPIP51 in DENV/ZIKV infection (Suppl. Fig. 3). Interestingly, viral infection induced a drastic relocalization of ER-resident VAPB into large NS3-positive foci reminiscent of CMs, suggesting that DENV and ZIKV alter VAPB/PTPIP51-dependent ERMC by physically sequestering VAPB. Altogether, these data strongly support that DENV and ZIKV target ERMC tethering complexes.

## DENV and ZIKV perturb mitochondrial respiration

The remodelling of organelles such as ER or mitochondria by flaviviruses and maintaining vROs is presumably highly energy-consuming while being a source of cellular stress. Considering that mitochondria is the powerhouse of the cell, we investigated the impact of flaviviral infection on the oxidative respiration process in mitochondria, which is the main source of energy in the form of adenosine triphosphate (ATP) in cells. Briefly, we analyzed mitochondrial respiration properties in uninfected-and infected-living Huh7.5 cells at 24, 48 and 72hrs post-infection by measuring the oxygen consumption rate (OCR) while sequentially adding oligomycin (ATP synthase inhibitor), FCCP (protonophore) and rotenone/antimycin A (complex I and III inhibitors, respectively) that directly act on the electron transport chain of the inner mitochondrial membrane (Fig. 3a). The OCR profile, an indicator of mitochondrial respiration, slightly increased at 24hrs post-infection with both viral infections compared to the uninfected condition (Fig. 3b, e). Very interestingly, the OCR profiles were shifted down at later time points of the infection at levels attributed to the non-mitochondrial oxygen consumption (Fig. 3c-e) suggesting that mitochondria are no longer able to efficiently perform oxidative respiration. Analyzing the OCR profiles revealed significant increases of the basal respiration, the ATP-linked respiration, and the maximal respiration at 24hrs post-infection whereas a drastic significant shut-off of these indicators was observed at 72hrs post-infection demonstrating a time-dependent modulation of the oxidative respiration by flaviviruses. (Fig. 3e).

**Figure 3:**
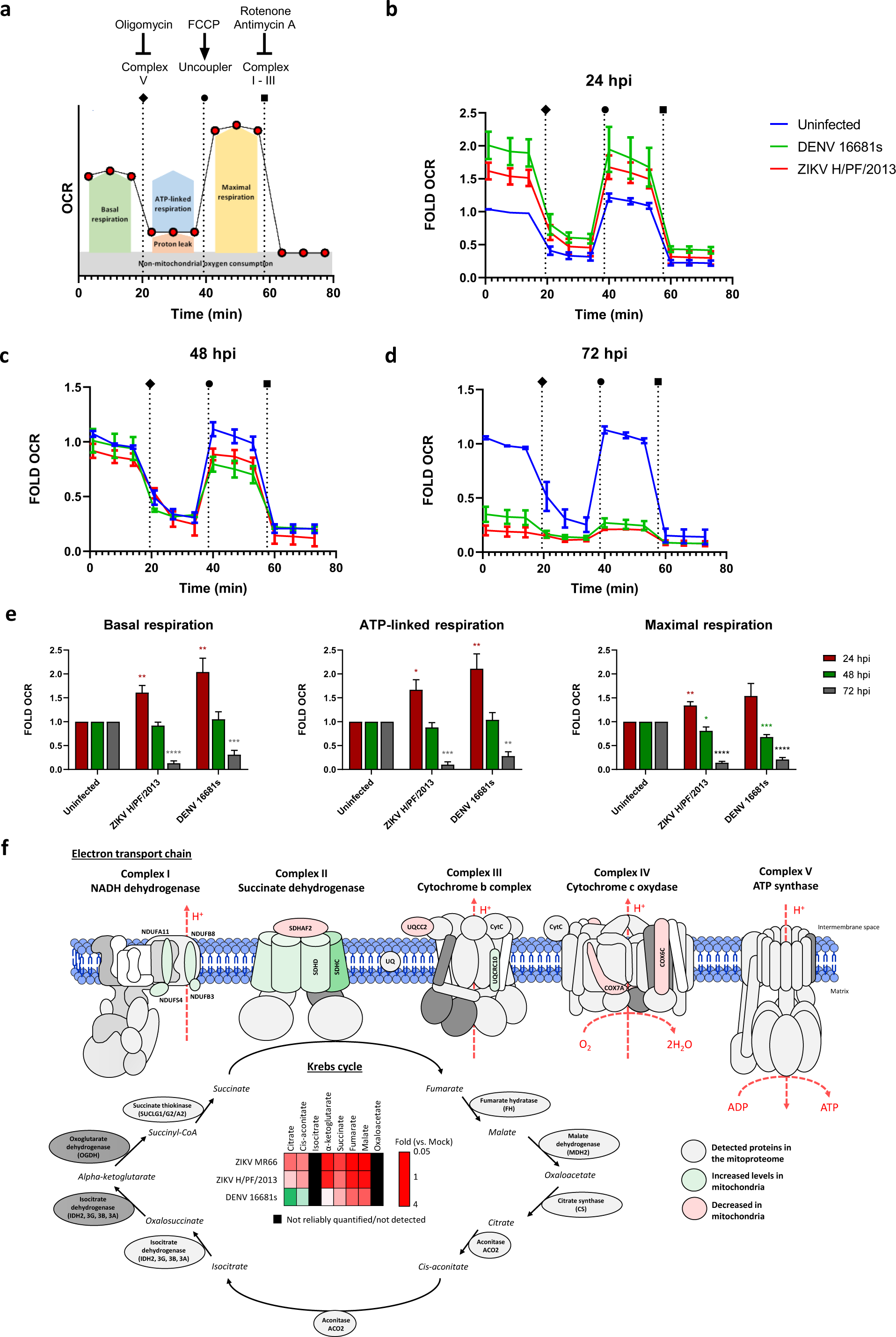
DENV and ZIKV perturb mitochondrial respiration. Huh7.5 cells were infected with DENV 16681s (MOI=1) or ZIKV H/PF/2013 (MOI=10), or left uninfected for 24, 48 and 72hr before analysis. (a) Schematic representation of the respiration profile in living cells generated with the Seahorse technology. The ATP synthase inhibitor oligomycin, the proton ionophore carbonyl cyanide p-trifluoromethoxyphenylhydrazone (FCCP), and a mixture of mitochondrial complex III inhibitor antimycin A / mitochondrial complex I inhibitor rotenone are sequentially added to analyze various parameters of the respiration. (b-d) The oxygen consumption rates (OCR) of uninfected and infected cells were analyzed at the indicated time points (N=5). (e) Based on the data of (3b-d), basal respiration, ATP production and maximal respiration were determined. (f) DENV/ZIKV-infected and control cells were analyzed two days post-infection for their mitochondrial protein composition following mass spectrometry on purified mitochondria (N=4), as well as for the abundance of the Krebs cycle metabolites measured by GC-MS in 4 independent experiments. Schematic representation of the mitochondrial electron transport chain coupled to the Krebs cycle is shown. Red elements indicate proteins/metabolites which were downregulated upon DENV/ZIKV infection. In contrast, green elements highlight upregulated proteins/metabolites. The expressions of the proteins in grey were not significantly impacted by flaviviral infections. All values were normalized to the uninfected condition. Isocitrate and oxaloacetate were not detected using GC-MS in these experiments.

To gain more insight about the causes of this respiration alteration at the late time points of flaviviral infection, we analyzed the mitochondrial levels of proteins involved in the Krebs cycle and the electron transport chain (ETC) by performing quantitative mass spectrometry on mitochondria purified from uninfected and ZIKV-and DENV-infected-Huh7.5 cells 48 hrs post-infection (Fig 3f). We have included in the analysis a second ZIKV strain from the African lineage, namely MR766. This mitoproteomic analysis identified changes in the stoichiometry of several proteins of the ETC proteins compared to uninfected cells (Fig. 3f, Suppl. Table 1, Suppl. Fig. 4a). Most notably, the composition of the complex II (succinate dehydrogenase), which is a component of both the Krebs cycle and the ETC was changed upon infection. Indeed, mitochondrial levels of sub-units D and C of SDH were increased upon DENV and ZIKV MR766 infections, respectively, compared to other subunits. Interestingly, the SDH assembly factor SDHAF2 was less abundant in mitochondria from DENV-infected cells and this phenotype was confirmed by western blotting on purified mitochondria (Suppl. Fig. 4b). However, in contrast to SDH, the mitochondrial levels of all other enzymes of the Krebs cycle remained unchanged (Fig 3f, Suppl. Table 1). The changes in SDH composition correlated with a drastic decrease of several metabolites of the Krebs cycle (i.e., α-ketoglutarate, succinate, fumarate and malate) as measured by gas chromatography-coupled mass spectrometry (GC-MS) (Fig. 3f, Suppl. Fig. 4c). Interestingly, in the case of DENV infection, the levels of citrate and cis-aconitate were specifically increased, which suggests a dysfunction of the Krebs cycle downstream aconitase activity. In addition to complex II, the stoichiometries of complexes I, III and IV were also altered in DENV-and ZIKV-infected cells with COX6C, COX7A2, COA4 and NDUFS4 being modulated by all three tested flaviviruses (Fig. 3f, Suppl. Fig. 4a, Suppl. Table 1). In contrast, some modulated ETC proteins were specific to either virus (Suppl. Fig. 4a). Flow cytometry analysis of infected cells with MitoTracker Orange showed that DENV and ZIKV did not induce a notable decrease in the mitochondrial potential at 48hrs post infection (Suppl. Fig 4d). This strongly supports that the observed respiration and mitoproteome phenotypes were not the result of a loss of mitochondrial integrity at that time point. Altogether, these data combining respirometric, proteomic and metabolomic approaches clearly demonstrate that ZIKV and DENV interfere with the mitochondrial respiratory metabolism.

**Figure 4:**
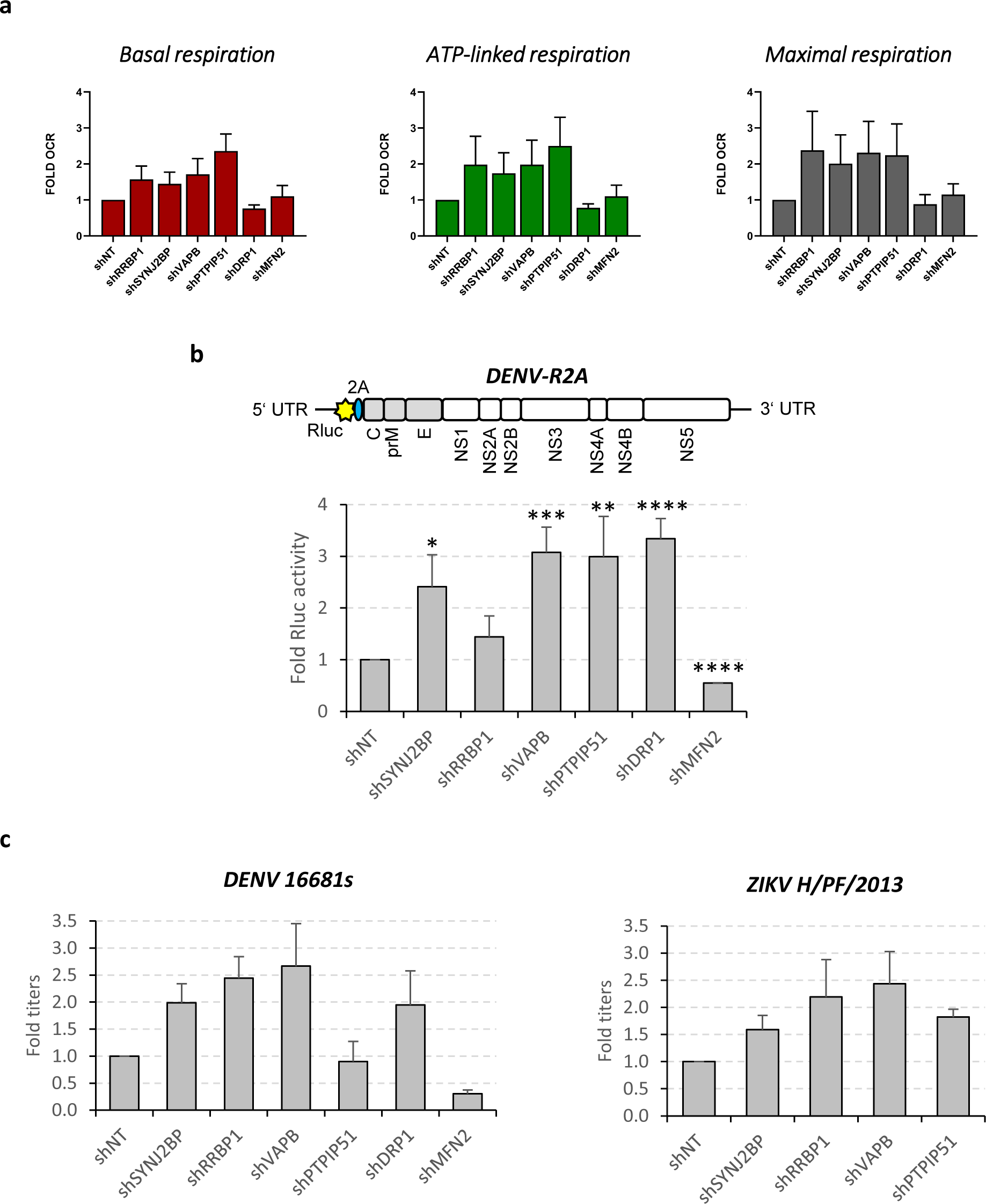
ERMC proteins negatively regulate mitochondrial respiration and viral replication. (a) Huh7.5 cells were transduced with lentiviruses expressing shRNAs which target the indicated proteins (MOI=4). Four days later, cells were trypsinized, counted and processed for measurements of various parameters of the oxygen consumption rate using the Seahorse technology as described in the Material and Methods section. (N=6). Data were normalized to the non-target shRNA (shNT) control condition. (b) Cells were transduced as in (a). Two days post transduction, cells were infected with DENV-R2A reporter viruses which express *Renilla reniformis* luciferase (Rluc) at a MOI of 0.01. Two days post-infection, the luciferase activity was measured as a read-out of viral replication and normalized to the shNT control condition. (c) Two days post-transduction, cells were infected with wildtype DENV 16681s or ZIKV H/PF/2013 (MOI=0.1). Two days post-infection, the infectious titers of extracellular viral particles were determined by plaque assays. All values were normalised to the shNT condition.

## ERMC alteration increases mitochondrial respiration

Since DENV and ZIKV infections induce a shut-off of the mitochondrial oxygen consumption over time and disrupt ERMCs, we investigated the impact of this sub-cellular compartment alteration on respiration. To that aim, we analyzed the oxygen consumption of living cells following expression knockdown of ERMC tethering proteins by transducing cells with shRNA-expressing lentiviruses. Indeed, it is well established that knockdown of ERMC proteins destabilizes physical contacts between ER and mitochondria organelles (Anastasia *et al*, 2021; Deng *et al*, 2008; Stoica *et al*, 2014; Duan *et al*, 2018). We identified shRNAs whose expression led to efficient silencing of SYNJ2BP, RRBP1, VAPB and PTPIP51 without reducing cellular viability 4 days post-transduction (Suppl. Fig. 5a-b). The IP3R1-GRP75-VDAC1 tethering complex was voluntarily omitted from the analysis because we reasoned that knocking these factors down may induce pleiotropic effects related to their ion channel activity and their role in calcium homeostasis and not to the contacts *per se* (Hartmann & Verkhratsky, 1998). To exclude that potential phenotypes would be due to changes in mitochondrial morphodynamics (which can influence viral replication), we used confocal microscopy to control that ERMC protein knockdown did not change the morphology of the mitochondrial network compared to the non-target shRNA (shNT) control condition (Suppl. Fig. 5d). As control, knockdown of fission factor DRP1 and fusion factor MFN2 led to mitochondrial elongation and fragmentation, respectively (Suppl. Fig. 5a, d). Moreover, flow cytometry analysis of ERMC protein-depleted cells showed no impact of ERMC alteration on the mitochondrial potential (Suppl. Fig. 5c). Interestingly, the mitochondrial oxygen consumption in Huh7.5 cells in which ERMC proteins were knocked down generally increased as compared to the shNT control condition (Fig. 4a). In contrast, no obvious changes in the OCR parameters were observed when mitochondrial morphodynamics was modulated upon DRP1 and MFN2 knockdown. These data show that ERMC alteration by DENV and ZIKV stimulates respiration independently of mitochondria morphodynamics and may explain why respiration is increased at early time points. Such a viral regulation also might attenuate the respiratory stress induced later in the infection.

## ERMC protein knockdown increases viral replication

We next investigated the impact of RNAi-mediated alteration of ERMC on viral replication. Huh7.5 cells in which ERMC proteins were knocked down were infected with a DENV reporter virus (DENV-R2A) expressing *Renilla reniformis* luciferase (Rluc), allowing us to evaluate DENV replication levels by measuring bioluminescence in cells at 2 days post-infection. We observed an increase in DENV-R2A replication upon ERMC protein knockdown, which was most pronounced when VAPB and PTPIP51 were depleted (Fig. 4b). Consistently, comparable phenotypes were obtained when cells were infected with either wild-type DENV2 16681s or ZIKV H/PH/2013 since the production of infectious viral particles was increased upon knockdown of ERMC proteins as measured by plaque assays (Fig. 4c). As controls and in line with previous observations (Chatel-Chaix *et al*., 2016), the modulation of mitochondrial morphodynamics showed the expected phenotypes. Indeed, the silencing of DRP1, which results in enhanced mitochondrial elongation, stimulated DENV replication while reduction of MFN2 expression impaired it (Fig. 4b-c). Overall, these data indicate that ERMC proteins restrict viral replication and further support that the DENV-and ZIKV-induced modulation of the physical contacts between mitochondria and ER is proviral.

## NS4B viral protein inhibits the mitochondrial respiratory metabolism

The conserved flaviviral NS4B protein is essential for viral replication and is a component of vROs (Chatel-Chaix *et al*; Grant *et al*, 2011; Kelly *et al*, 2010; Welsch *et al*., 2009; Zou *et al*, 2014). Since it was shown that DENV NS4B expression induces mitochondria elongation (Chatel-Chaix *et al*., 2016), we hypothesized that this viral protein was at least in part responsible for the observed alteration of mitochondrial respiration in DENV-and ZIKV-infected cells. First, we showed that the transient expression of ZIKV NS4A-2K-NS4B precursor and 2K-NS4B pseudo-mature proteins (2K serving as a signal peptide for NS4B) in Huh7.5-T7 cells induced mitochondria elongation as DENV NS4B, suggesting that ZIKV and DENV NS4B proteins modulate mitochondria functions via similar molecular mechanisms. (Suppl. Fig. 6a-b).

We next investigated the impact of NS4B expression on mitochondrial respiratory functions. To ensure that all cells stably expressed this viral protein, we transduced Huh7.5 cells using lentiviruses expressing ZIKV or DENV HA-tagged NS4B and conferring resistance to puromycin. Following selection with this antibiotic, we verified the proper expression of NS4B by western blotting (Fig. 5a) and subsequently measured the abundance of the Krebs cycle metabolites by GC-MS. As in infected cells, the overexpression of both DENV and ZIKV NS4B significantly reduced the levels of α-ketoglutarate, succinate and fumarate (Fig 5b). Consistently, this correlated with a decrease in basal respiration, ATP-linked respiration and maximal respiration (Fig. 5c) which was more pronounced for ZIKV NS4B-HA. Comparable phenotypes on oxygen consumption rates were observed when DENV and ZIKV NS4B proteins were expressed as NS4A-2K-NS4B precursors (Suppl. Fig 6c-d). Altogether, these data demonstrate that NS4B is one of the main viral determinants contributing to the decrease of the oxidative respiration observed at the late time points of DENV and ZIKV infections.

**Figure 5:**
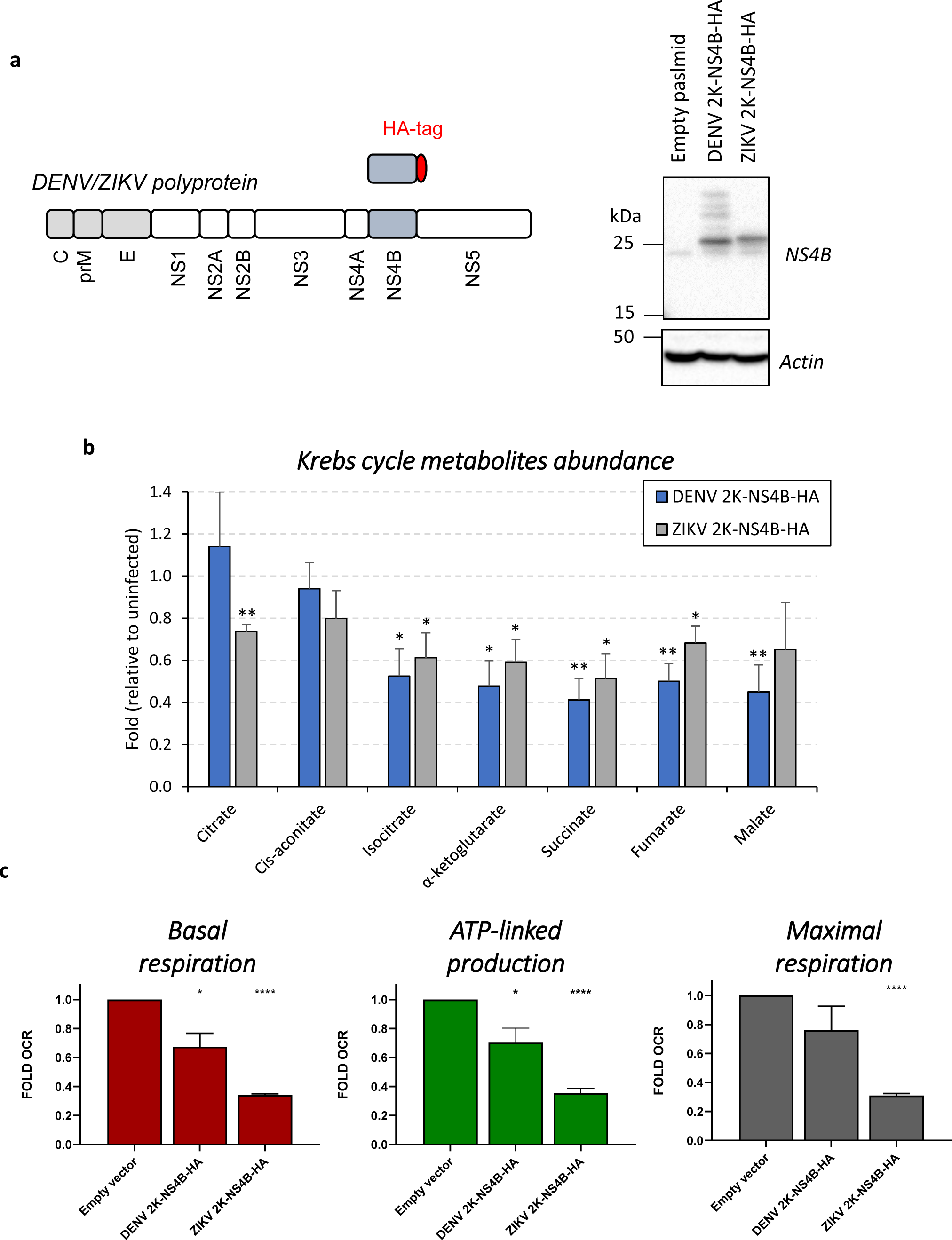
NS4B viral protein inhibit the mitochondrial respiratory metabolism. Huh7.5 cells were transduced with lentiviruses expressing HA-tagged DENV or ZIKV NS4B proteins including the N-terminal 2K signal peptide (MOI= 2-4). Transduced cells were selected with puromycin and analyzed 4 days post-transduction for: (a) NS4B expression using western blotting, (b) the abundance of Krebs cycle metabolites using GC-MS in 3 independent experiments, and (c) their basal respiration, ATP production and maximal respiration using the Seahorse technology (N=3). All values were normalised to the empty vector control condition. For (b) and (c) an equal number of cells for all conditions was used.

**Figure 6:**
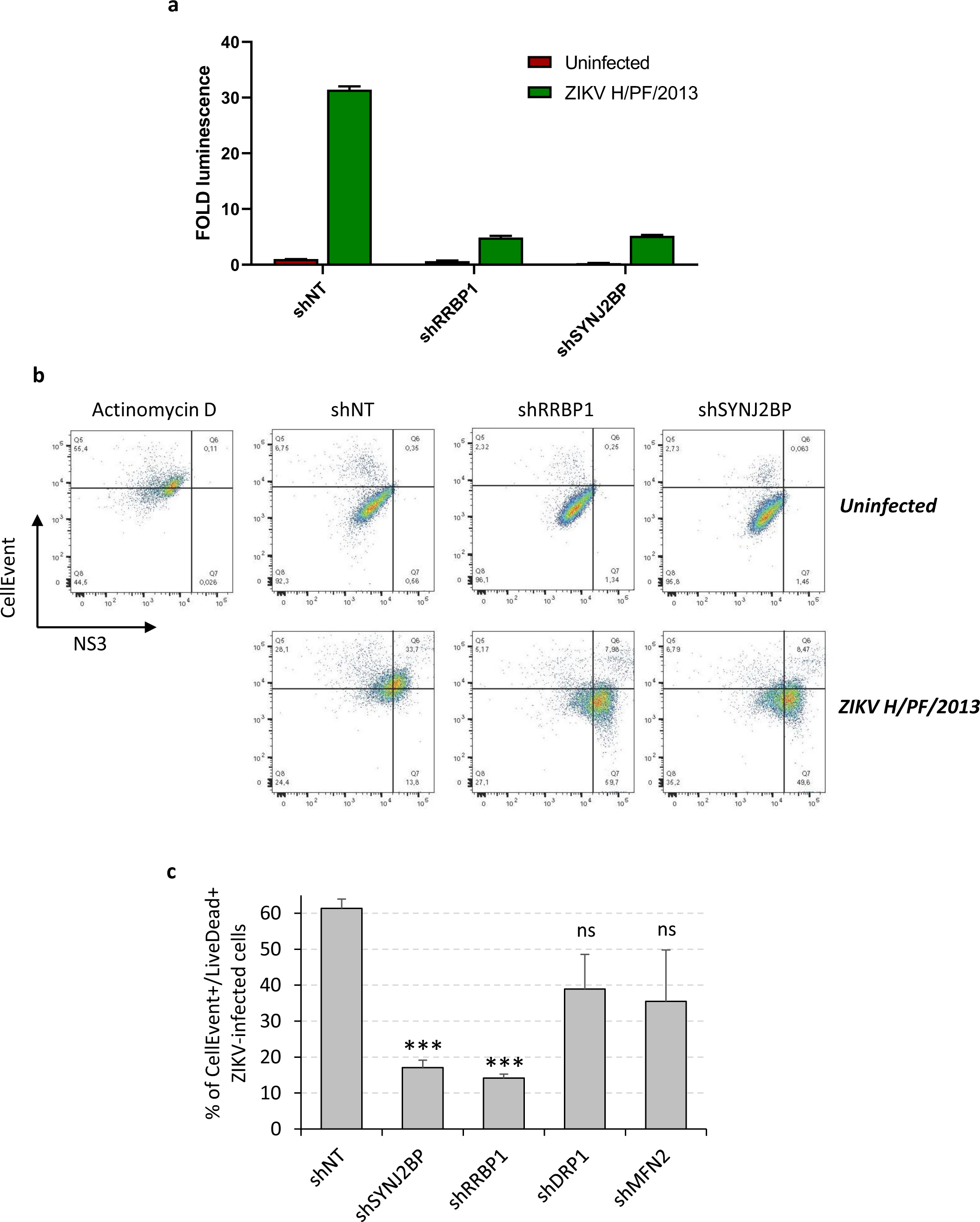
ERMC alteration dampens ZIKV-induced apoptosis. (a) Huh7.5 cells were transduced with lentiviruses expressing shRNAs which target the indicated proteins (MOI=4). Two days post-transduction, cells were infected with ZIKV H/PF/2013 (MOI=20) or left uninfected. Two days post-infection, cell lysates were prepared, and apoptosis induction was measured by measuring bioluminescence using the Caspase-Glo 3/7 assay kit. (b) Cells were transduced and infected as in (a). Two days post-infection (*i.e.*, 4 days post-transduction), apoptosis induction, cell death and ZIKV infection were detected by flow cytometry using the CellEvent caspase-3/7 green reagent, the amine reactive viability dye LIVE/DEAD aqua fixable stain, and anti-NS3 antibodies, respectively. Actinomycin D treatment (2.5 µM, 24 hours) was used as a positive control of apoptosis induction. The plots show representative results of the caspase activity in function of NS3 expression. (c) Gated ZIKV-infected cells (*i.e.*, NS3+ cells) were analyzed for apoptosis induction and cell viability using the CellEvent caspase-3/7 green and LIVE/DEAD biosensors, respectively (N=3). All values were normalised to the shNT condition.

## RRBP1 and SYNJ2BP regulate ZIKV-induced apoptosis

We have previously shown that CM integrity relies on VCP ATPase activity which is required to dampen ZIKV-induced apoptosis (Anton *et al*., 2021). Since CMs are physically connected to mitochondria through residual associated ER membranes and considering that ERMC were reported to regulate mitochondria-dependent cell death (Carpio *et al*, 2021; Giamogante *et al*, 2021; Suresh, 2019; Verfaillie *et al*, 2012), we hypothesized that the alteration of ERMC integrity dampens apoptosis during flaviviral infection. To test this, we measured the activity of caspase 3 (the major regulator of the execution phase of apoptosis) in Huh7.5 cells upon RRBP1 or SYNJ2BP silencing. We chose this ERMC tethering couple because it was the one that was the most impacted by ZIKV infection (Fig. 2b-c). Very interestingly, upon RRBP1 or SYNJ2BP depletion, the activity of caspase 3/7 was reduced in lysates of ZIKV-infected cells as measured using a bioluminescence-based assay (Fig. 6a). We confirmed these phenotypes by flow cytometry using the CellEvent caspase-3/7 green reagent, a fluorescent biosensor of caspase 3/7 activity, and anti-NS3 antibodies to detect ZIKV-infected cells (Fig. 6b). While 61.2% of the cells were apoptotic (*i.e.*, CellEvent-positive) in control infected conditions (shNT), this proportion was reduced to 13.2% and 15.3% when RRBP1 and SYNJ2BP were knocked down, respectively. Moreover, the proportion of ZIKV infected cells which were non-apoptotic (*i.e.*, NS3-positive/CellEvent-negative cells) drastically increased under these conditions (13.8% for shNT versus 59.7% and 49.6% for shRRBP1 and shSYNJ2BP, respectively). Furthermore, by combining these markers with the amine reactive viability dye LIVE/DEAD aqua fixable stain (Fig. 6c), we showed that the silencing of RRBP1 and SYNJ2BP decreased the proportion of dead ZIKV-infected cells, namely cells that were apoptotic and exhibited a ruptured plasma membrane (% of NS3-gated CellEvent-positive and LIVE/DEAD-positive cells). Finally, as expected, no apoptosis was observed in any uninfected conditions demonstrating that ERMC destabilization *per se* does not induce apoptosis (Fig. 6b). In contrast, as positive control, treatment of the cells with actinomycin D robustly induced apoptosis. Altogether, these data support that ZIKV alters ERMC integrity to dampen infection-induced apoptosis and promote replication.

## DISCUSSION

In this study, we demonstrate that DENV and ZIKV modulate the mitochondrial oxidative respiration and virus-induced apoptosis in favor of viral replication notably by altering the physical contacts between mitochondria and the ER. Our observations are consistent with the fact that ERMC were reported to play a crucial role in the induction of the apoptosis pathway, as explained below (Carpio *et al*., 2021; Giamogante *et al*., 2021; Suresh, 2019; Verfaillie *et al*., 2012). Such morphological remodeling allows flaviviruses to create a proviral cytoplasmic environment. Although there is reported evidence that mitochondrial fission is initiated at sites where the ER tubules wrap mitochondria and constrict mitochondria prior to DRP1 recruitment (Friedman *et al*., 2011), we did not observe any changes in mitochondria morphology upon ERMC protein knockdown in Huh7.5 cells (Suppl. Fig. 5d) supporting that the phenotypes reported here are independent of DRP1-mediated fission regulation by flaviviruses (Barbier *et al*., 2017; Chatel-Chaix *et al*., 2016).

Flaviviruses induce membranous replication factories from the ER membranes (Chatel-Chaix & Bartenschlager, 2014; Paul & Bartenschlager, 2015). More recently, it was shown that DENV and ZIKV induce the elongation of mitochondria in the vicinity of CMs (Barbier *et al*., 2017; Chatel-Chaix *et al*., 2016). As it was already observed (Chatel-Chaix *et al*., 2016), we noticed that mitochondria remain connected to CMs via residual associated ER membranes. This raises the hypothesis that the morphogenesis of CMs uses the reticulo-mitochondrial interface as a source of ER membranes via the destabilization of this specific cytoplasmic compartment. In addition, we show that DENV and ZIKV alter ERMC by decreasing the abundance of three different tethering protein complexes (RRBP1-SYNJ2BP, IP3R1-VDAC1 and VAPB-PTPIP51). We did not observe any difference in the expression level of VDAC1, VAPB and PTPIP51. Interestingly, RRBP1 levels were reduced after 48 hours of infection with both viruses while a ZIKV-specific SYNJ2BP product exhibiting a higher molecular weight was detected. Whether these results from a flavivirus-specific regulation of alternative splicing or of posttranslational modifications will be explored in future studies. While VAPB-PTPIP51 and IP3R1-VDAC1 were reported to contribute to calcium homeostasis, autophagy or in phospholipid transfer (Colombini, 2016; De Vos *et al*., 2012; Gomez-Suaga *et al*, 2017; Yeo *et al*, 2021), the contribution of the RRBP1-SYNJ2BP complex to specific cellular function beyond contributing to ERMCs is unknown. This could explain why DENV and ZIKV target specifically this protein-protein couple to separate ER and mitochondria without affecting essential cellular processes. Very interestingly, a recent mass spectrometry-based study has shown that in human fibroblasts, herpes simplex virus type 1, influenza A virus and betacoronavirus HCoV-OC43 decrease the abundance of mitochondrial proteins involved in ERMC whereas the human cytomegalovirus (HCMV) mostly positively regulates them (Cook *et al*, 2022). Increasing the abundance of ERMC proteins such as VAPB, MFN1/2 fusion proteins, fission-associated proteins DRP1 and MFF, and ER-mediated calcium transfer proteins PTPIP51 and VDAC1 occurred very early during the infection (8 hours post-infection). This HCMV-mediated regulation of ERMCs favors cellular processes such as ER-to-mitochondria calcium flux, mitochondrial fragmentation, and reshaping of cristae for the benefit of HCMV viral particle production. This highlights that viruses from other families than *Flaviviridae* have evolved to co-opt mitochondrial functions via diverse strategies.

The biogenesis of vROs occurs early in infection and their maintenance presumably requires high energy needs in the form of ATP. Consistently, we show here that in the first 24 hours of infection, DENV and ZIKV significantly increase the oxidative respiration followed by a near complete shut-down of this bioenergetic process later in the infection, which correlated with a decrease in the levels of several metabolites of the Krebs cycle (Fig. 3). This might result from the mitochondrial stress and the onset of cytopathic effects generated by the infection. The close analysis of oxygen consumption rate profiles using the Seahorse technology with different inhibitors of the ETC revealed that global respiration was impacted by DENV and ZIKV rather than for instance only the maximal respiration. In line with our data, a recent study showed increases in oxygen consumption and ATP production in astrocytes at the early time points of the infection (*i.e.*, 18 to 24 hrs post-infection) (Ledur *et al*, 2020). In contrast, later in the infection, the phenotype was reversed with an observed decrease of the oxygen consumption at 48 hrs post-infection. These data support our model of high energy needs when DENV and ZIKV establish the infection notably during the biogenesis of vROs. It is worth mentioning that some studies reported contrasting phenotypes. For instance, it was reported that DENV infection increases oxidative respiration in Huh7 and HepG2 liver cells at 2 days post-infection (Barbier *et al*., 2017; El-Bacha *et al*, 2007). Another study showed an inhibition of the basal and maximal oxygen consumption in first-trimester primary trophoblasts infected with ZIKV, that correlated with the fragmentation of the mitochondrial network (Chen *et al*, 2020). In neurons, ZIKV infection also induced mitochondrial fission by decreasing the expression of the fusion factor MFN2 despite low levels of viral replication (Yang *et al*, 2020). Pharmacological inhibition of DRP1 (and hence, of fission) reduced ZIKV-induced cell death in this model. This suggests a functional interplay between mitochondrial morphodynamics, the respiratory metabolism and virus-induced apoptosis, which dictates the fate of the infected cell and would be regulated by flaviviruses. While some of these results may seem contradictory and might be explained by the used viral strains and/or cell types more prone to cell death induction (and metabolism suppression) upon infection/stress (*e.g.*, neurons, immune competent cells), we believe that our kinetic analysis described here showing that the time of infection (and most probably the extent of viral protein and remodeled organelles accumulation) is an important determinant of the flaviviral impact on the respiratory metabolism and partly explains these discrepancies. Furthermore, we used here the Huh7.5 cell line which is deficient for the RIG-I-dependent signaling and interferon production. Thus, we can rule out that the observed phenotypes are due to the activation of antiviral innate immunity. The fact that ERMC protein knockdown increased the oxygen consumption rate (Fig. 4a) supports the model in which DENV and ZIKV alter ERMCs to stimulate mitochondrial activity to favor vRO formation and delay cell death at the late time points of infection. Very interestingly, we demonstrate that NS4B viral protein is at least partly responsible for the observed decreases in the oxygen consumption rate and the levels of Krebs cycle metabolites. While it remains unknown how NS4B can interfere with mitochondrial respiration, it is noteworthy that this protein is absolutely required for viral RNA replication and is the target of highly potent direct-acting antivirals including one currently challenged in clinical trial (Chatel-Chaix *et al*., 2015; Kaptein *et al*, 2021; Moquin *et al*, 2021; van Cleef *et al*, 2013; Xie *et al*, 2011; Zou *et al*, 2015; Zou *et al*., 2014).

Apoptosis is a programmed cell death that is induced by both extrinsic and intrinsic pathways. The intrinsic pathway relies on the activation of the caspase 3 via the release of BCL-2 family proteins from mitochondria (Elmore, 2007). Given that ERMC were reported to regulate this cell death pathway, we investigated whether their alteration could influence ZIKV-induced apoptosis. In this study, we demonstrate that the alteration of ERMCs by knocking down RRBP1 and SYN2BP decreases apoptosis in ZIKV-infected cells. ERMCs notably regulate this cell death induction pathway through IP3R1 and VDAC1 which interact and act as calcium channels to directly transfer calcium from the ER to mitochondria. Since an overload of calcium in mitochondria leads to apoptosis (Orrenius *et al*, 2003), it is tempting to speculate that the loss of ERMCs reduces the amounts of mitochondrial calcium and thus, delays the induction of apoptosis. Mitochondria fission positively regulates the activation of apoptosis, notably through the interaction between DRP1 and BAX, a pro-apoptotic factor (Jenner *et al*, 2022; Montessuit *et al*, 2010; Oettinghaus *et al*, 2016; Park *et al*, 2015). However, RRBP1 and SYN2JBP knockdown did not result in apparent changes in mitochondria morphology, thus allowing us to conclude that these phenotypes were not related to the mitochondria elongation induced by DENV and ZIKV. Interestingly, we have already showed that the integrity of CMs during ZIKV infection is required to both dampen apoptosis and maintain the elongated morphology of mitochondria (Anton *et al*., 2021). This suggests that both DENV/ZIKV-induced mitochondria elongation and ERMC alteration independently regulate apoptosis. It is interesting to note that in our cell culture system, mitochondria integrity appeared to be preserved upon DENV/ZIKV infection and ERMC protein knockdown since we did not detect any notable changes in the mitochondrial membrane potential (Suppl. Fig. 4d, 5c), implying that these cells are quite resistant to the cytopathic effects induced by mitochondrial stress.

Finally, the ERMC compartment also serves as a signalling platform during early antiviral innate immunity leading to the induction of type I and III interferons (IFN). When RIG-I senses a foreign RNA, it is translocated to the surface of the mitochondria and binds mitochondrial antiviral-signalling protein adaptor (MAVS) which is located at the ERMC (Horner *et al*., 2011). Several inhibition mechanisms of RIG-I activation by flaviviruses have been discovered in the last decade (Chatel-Chaix *et al*., 2016; Gack & Diamond, 2016; Gack *et al*, 2007; Liu *et al*, 2012; Manokaran *et al*, 2015; Riedl *et al*, 2019; Serman & Gack, 2019; Tremblay *et al*, 2019). Thus, perturbing the reticulo-mitochondrial interface may constitute another strategy to inhibit or delay cellular processes that could be harmful for viruses. While we can rule out that our phenotypes in RIG-I-deficient Huh7.5 cells are due to interferon induction, it will be interesting to evaluate in the future whether the alteration of the reticulo-mitochondrial interface by flaviviruses contributes to countering antiviral immunity in immune-competent cells.

Overall, our data demonstrate that the morphological perturbations of the reticulo-mitochondrial interface by DENV and ZIKV modulate mitochondrial respiratory metabolism to sustain the energetic needs of flaviviral life cycle. This study further supports a model in which these viruses perturb ERMC to hijack specific host factors that are required for CM morphogenesis, and vice-versa. More studies are required to better understand these mechanisms at a molecular level, including the specific viral determinants involved in the alteration of ERMCs and the mitochondrial functions regulated by this sub-cellular compartment. It will be also interesting to evaluate how this impacts ERMC function other than apoptosis and innate immunity, such as calcium homeostasis or lipid metabolism.

## MATERIAL AND METHODS

### Cell lines and virus strains

Huh7.5, HEK293T, VeroE6, and HeLa cells were cultured in Dulbecco’s modified Eagle medium (DMEM, Life Technologies) supplemented with 10% fetal bovine serum (Wisent), 1% non-essential amino acids (Life Technologies) and 1% penicillin-streptomycin (Life Technologies). The generation of the Huh7.5-T7 cell line expressing the T7 RNA polymerase was reported elsewhere (Anton *et al*., 2021). These cells were cultured in the presence of 5 µg/mL blasticidin (Thermo-Fisher). Huh7.5 cells stably expressing mito-mTurquoise2 were produced exactly as before using lentiviral transduction (Chatel-Chaix *et al*., 2016) and were maintained in 5 µg/mL zeocin (Life Technologies). ZIKV H/PF/2013 and ZIKV MR766 strains were provided by the European Virus Archive Global (EVAg). Virus stocks were generated by amplification in VeroE6 cells following inoculation with a multiplicity of infection (MOI) of 0.001. Virus aliquots were stored at 80°C until use. Infectious titers were determined by plaque assays exactly as reported before (Freppel *et al*, 2018). DENV2 16681s and reporter Rluc-expressing DENV-R2A particles were generated using a reverse genetics system (a kind gift of Ralf Bartenschlager) (Fischl & Bartenschlager, 2013) and by electroporating VeroE6 cells with *in vitro*-transcribed DENV RNA genomes as reported before (Mazeaud *et al*, 2021).

### Antibodies

Rabbit anti-DENV NS4B (GTX124250; cross-reactive for ZIKV), rabbit anti-ZIKV NS4B (GTX133311), rabbit anti-ZIKV NS3 (GTX133309) and mouse monoclonal anti-DENV NS3 (GTX629477; cross-reactive for ZIKV) were obtained from Genetex. Rat polyclonal antibodies against DENV2 16681 NS3 which are cross-reactive with ZIKV NS3 were generated by Medimabs (Montreal, Canada) as reported before (Anton *et al*., 2021). Mouse anti-VDAC1 (ab14734) and mouse anti-GRP75 (ab2799) were obtained from Abcam. Rabbit anti-IP3R1 (NBP2-22458) antibodies were obtained from Novus Biologicals. Mouse anti-VAPB (66191-1-Ig) was obtained from ProteinTech. Mouse anti-RRBP1 (MA5-18302) was obtained from Invitrogen. Rabbit anti-PTPIP51 (HPA009975) and rabbit anti-SYNJ2BP (HPA000866) were obtained from Sigma-Millipore.

### DNA cloning

The cloning of the constructs expressing DENV/ZIKV NS4B proteins in a T7 RNA polymerase-dependent manner was previously described (Anton *et al*., 2021; Chatel-Chaix *et al*., 2015). For generating the lentiviral plasmids expressing HA-tagged 2K-NS4B or NS4A-2K-NS4B proteins, corresponding coding sequences were amplified using the plasmids above as templates (Fischl & Bartenschlager, 2013). DNA fragments were cloned into the AscI/SpeI or BamHI/SpeI cassettes of the pWPI lentiviral plasmid for ZIKV and DENV proteins, respectively (Chatel-Chaix *et al*., 2016).

### Lentivirus production, titration, and transduction

Knockdowns of ERMC proteins were achieved by transduction with lentiviruses expressing MISSION shRNA from MilliporeSigma (shSYNJ2BP: TRCN0000121988; shRRBP1: TRCN0000117408; shVAPB: TRCN0000152520; shPTPIP51: TRCN0000135580). Constructs expressing shDRP1 and shMFN2 were already described (Chatel-Chaix *et al*., 2016). For lentivirus production, HEK293T cells were transfected with packaging plasmids pCMV-Gag-Pol, pMD2-VSV-G and pLKO-shRNA or pWPI expressing NS4B using 25 kDa linear polyethylenimine (Polysciences Inc.) exactly as before (Anton *et al*., 2021; Chatel-Chaix *et al*., 2016; Mazeaud *et al*., 2021). Two days post-transfection, lentivirus-containing medium was collected, filtered and stored at −80°C until use. Lentivirus titration was performed in HeLa cells. Cells were seeded at 50,000 cells/well in 24-well plates and lentivirus-containing medium was titrated in 10-fold serial dilution 10^-1^ to 10^-3^ in duplicate. Transduced cells were selected by antibiotic selection one-day post-transduction treatment with 1 µg/mL puromycin. Six days post-transduction, cells were washed once in PBS, and then fixed and stained with 1% crystal violet/10% ethanol for 20 minutes. Stained cells were washed with tap water. Colony-forming unit were counted and titers were determined taking into consideration the dilution factor. Huh7.5 cells were transduced with lentiviruses at a MOI of 5-10 in the presence of 8 µg/mL polybrene.

### Cell viability assays

7,500 Huh7.5 cells/well in 100µl-DMEM were seeded in 96-well plates and transduced as indicated above. Four days post transduction, 20 µL of 3-(4,5-dimethylthiazol-2-yl)-2,5-diphenyltetrazolium bromide (MTT) at 5 mg/mL were added to the medium for 1 to 4 hr at 37°C. Medium was removed and 150 µL of 2% (v/v) of 0.1 M glycine in DMSO (pH 11) were added to dissolve the MTT precipitates. Absorbance at 570 nm was read with Spark multimode microplate reader (Tecan).

### Renilla luciferase assay

1.10^5^ Huh7.5 cells/well were plated in 12-well plates in triplicates and transduced as indicated above. The day after, the culture medium was replaced. Two days post-transduction, cells were infected with virus DENV-R2A reporter virus (MOI∼0.01). 2 days later, the medium was removed, and cells were lysed in 200 µl of luciferase lysis buffer (1% Triton-X-100; 25 mM Glycyl-Glycine, pH7.8; 15 mM MgSO4; 4 mM EGTA; 1 mM DTT added directly prior to use). 30 µl of lysates were transferred into a white 96-well plate. Luminescence was read with a Spark multimode microplate reader (Tecan) after injection of 150 µl of assay buffer (25 mM Glycyl-Glycine pH7.8; 15 mM KPO_4_ buffer pH7.8; 15 mM MgSO_4_; 4 mM EGTA; 1 mM coelenterazine freshly added before the assay). All values were normalized to the shNT control condition.

### Virus production assays

2.10^5^ Huh7.5 cells/well were plated in 6-well plates and transduced in the presence of 8 µg/mL polybrene. The day after, culture medium was changed. Two days post-transduction, cells were infected with DENV2 16681s or ZIKV H/PF/2013 (MOI = 0.1). Three hours later, culture medium was changed. Two days post-infection, cell supernatants were collected, filtered at 0.45 µm, and kept at −80°C until use. 2.10^5^ VeroE6 cells/well were seeded in 24-well plates. The day after, cells were infected in 10-fold serial virus dilutions (10^-1^ to 10^-6^) in duplicate in complete DMEM. Three hours post-infection, the medium was removed, and replaced with serum-free MEM (Life Technologies) containing 1.5% carboxymethylcellulose (MilliporeSigma). Five (ZIKV) or seven (DENV) days post-infection, cells were fixed for 2 hours in 5% formaldehyde. Cells were washed vigorously with tap water and stained with 1% crystal violet/10% ethanol for 20 minutes. Cells were washed with tap water. Plaques were counted, and infectious titers in particles forming unit (PFU/mL) were calculated.

### Immunofluorescence-based confocal microscopy

For immunofluorescence of transiently expressed NS4B proteins, 50,000 Huh7.5-T7 cells/well in 24-well plates were seeded on sterile glass coverslips. The next day, cells were transfected with0.5 μg of NS4B-encoding plasmids using TransIT-LT1 Transfection Reagent (Mirus) according to the manufacturer’s instructions. Four hours post-transfection, culture media were changed. Eighteen hours post transfection, cells were rinsed twice in PBS and fixed in PBS containing 4% paraformaldehyde for 20 minutes.

For confocal microscopy of infected cells, Huh7.5 cells were seeded on coverslips before immunofluorescence assay and infected the day after with DENV2 16681s (MOI = 1-5) or ZIKV H/PF/2013 (MOI=5-10) 2-or 3-days post-infection, cells were washed three times with PBS and fixed for 20 minutes with PBS containing 4% PFA. Coverslips were rinsed 3 times with PBS and kept at 4°C in PBS until use. Prior immunostaining, cells were permeabilized for 15 min in PBS containing 0.2% Triton X-100, and then blocked for 1hr with PBS containing 5% bovine serum albumin (BSA) and 10% goat serum (Thermo-Fisher). Coverslips were incubated 2hrs with primary antibodies at room temperature diluted in PBS/5% BSA. Coverslips were washed three times in PBS and incubated for 1hr at room temperature and in the dark with Alexa Fluor (488, 568 or 647)-conjugated secondary antibodies (Life Technologies) diluted in PBS/5% BSA. Coverslips were washed three times in PBS and incubated 15 min in PBS containing 4’, 6’-diamidino-2-phenylindole (DAPI; Life Technologies) diluted 1/10,000 for nuclei staining. Coverslips were washed three times in PBS, and once in water before being mounted on slides with FluoromountG (Southern Biotechnology Associates). Cells were observed and imaged using a LSM780 confocal microscope (Carl Zeiss Microimaging) at the Confocal Microscopy Core Facility of the INRS-Centre Armand-Frappier Santé Biotechnologie. Images were processed with the Fiji software.

### Proximity ligation assays

Huh7.5 cells were cultured on coverslips and infected as above. Coverslips were washed three times with PBS and fixed for 20 min with PBS/4% paraformaldehyde. Coverslips were rinsed 3 times with PBS and kept at 4 °C in PBS until use. Prior to the assay, cells were permeabilized with 0.2% Triton X-100 for 15 min. Proximity ligation assays were performed using the Duolink PLA Kit (Millipore-Sigma) according to the manufacturer’s protocol. Briefly, cells were blocked with the kit buffer for 1 hr at 37°C in prior incubation with the indicated mouse and rabbit primary antibodies for 2 hours at room temperature. Coverslips were washed three times with the buffer A from the kit and incubated for 1 hr at 37 °C with PLUS and MINUS PLA probes. Coverslips were washed three times in buffer A and incubated for 30 min at 37°C with the ligation solution. Coverslips were washed three times with buffer A and incubated 100 min at 37°C with the kit amplification solution. After final washes with buffer B, coverslips were stained as above with rat anti-NS3 antibodies and Alexa Fluor488 anti-rat antibodies (see immunofluorescence-base confocal microscopy). Cells were then incubated for 15 min in 0.01% buffer B and a 1/10,000 dilution of DAPI (Life Technologies) for nuclei staining. Coverslips were washed three times in buffer B, then once in 0.01% buffer B and mounted on slides with FluoromountG (Southern Biotechnology Associates). Cells were observed and imaged using a LSM780 confocal microscope (Carl Zeiss Microimaging) at the Confocal Microscopy Core Facility of the INRS-Centre Armand-Frappier Santé Biotechnologie. The quantifications of the intracellular PLA dots were performed with the Fiji software. Briefly, 8-bit format images were processed for each channel. We increased the signal for NS3 to saturation in order to delineate the entire cell area. The threshold of PLA signal was adjusted to eliminate the non-specific background. A given PLA signal was considered positive when bigger than 0.02µm^2^ and having a circularity between 0.02 and 1.00. PLA dots were counted in each cell. Same acquisition and counting settings were applied to all data for each experiment.

### Transmission electron microscopy

Infected cells were prepared for transmission electron microscopy exactly as previously reported (Anton *et al*., 2021; Mazeaud *et al*., 2021). Briefly, Huh7.5 cells were seeded on Lab-tech chambers Slide^TM^ (Thermo Fisher) and infected with DENV 16681s MOI 1 or ZIKV H/PF/2013 MOI of 10. Two days later, cells were fixed in 2.5% glutaraldehyde in 0.1M sodium cacodylate buffer pH 7.4 overnight at 4°C and washed three times with washing buffer. Cells were postfixed for 1 hour with the washing buffer containing 1% aqueous OsO_4_ and 1.5% aqueous potassium ferrocyanide. Following three washes in washing buffer, samples were dehydrated in sequential dipping into a series of ethanol-dH_2_O solutions of increasing concentration up to 70% followed by a 1 hour-long staining with 2% uranyl acetate in 70% ethanol. Samples were washed twice in 70% ethanol and subjected to a progressive dehydration with up to 100% ethanol. A Graded Epon-ethanol series (1:1, 3:1) was used to infiltrate the samples before embedding in 100% Epon and polymerizing in an oven at 60 °C for 48hrs. A Diatome diamond knife using a Leica Microsystems EM UC7 ultramicrotome was used to cut serial sections (90-100 nm thick) from the polymerized blocks. Ultrathin sections were then transferred into 200-mesh copper grids and stained during 6 minutes with 4% uranyl acetate and 5 minutes in Reynold’s lead. Image acquisitions of TEM grids were performed with a FEI Tecnai G2 Spirit 120 kV TEM equipped with a Gatan Ultrascan 4000 CCD Camera Model 895 (Gatan, Pleasanton, CA) located at McGill University Facility for Electron Microscopy Research. TEM images were analyzed using the Fiji software for measurements of mitochondria perimeter and ERMC length. Ultrastructures were considered as ERMCs when the measured distance between ER and mitochondria was below 50nm. The ratio between these two values indicated the percentage of the mitochondrial perimeter in contact with ER.

### Oxygen consumption rate measurements

The day prior the assay, Seahorse 96-well sensor cartridges (Agilent) were hydrated overnight at 37°C in a non-CO_2_ incubator with 200 µl/well of the Seahorse XF calibrant (Agilent). The day of the assay, living cells were trypsinized and counted with Trypan blue. 50,000 cells/well were seeded into a Seahorse XF 96-well cell culture microplate (Agilent) in DMEM and incubated for at least 4 hours at 37 °C with 5% CO_2_. Culture medium was changed for 180 µl of Seahorse XF DMEM which was supplemented with 1 mM pyruvate, 2 mM glutamine and 10 mM glucose. Cells were incubated for 1 hour in a non-CO_2_ incubator at 37 °C. For measurements of the oxygen consumption rates, we used the Seahorse XF Cell Mito Stress Test kit (Agilent) according to the manufacturer’s instructions. Briefly, 20 µl oligomycin 10 µM, 22 µl FCCP 10 µM and 25 µl rotenone/antimycin A 5 µM were loaded into the ports of the sensor cartridges. Sensor cartridges were placed on top of the Seahorse XF cell culture microplate. Sequential drug addition (10-fold dilution) and time-lapse oxygen consumption rate measurements were achieved with a Seahorse XFe96 analyzer (Agilent). Data were analyzed using Wave 2.6.1 software to determine the basal respiration, the maximal respiration and the ATP production in each sample.

### Mitochondria affinity purification and quantitative LC-MS/MS

For the determination of mitochondrial proteome, four independent affinity purifications were performed for each experimental condition as follows. Huh7.5 cells were infected with DENV2 16681s (MOI=2), ZIKV H/PF/2013 (MOI=10) or ZIKV MR766 (MOI=2) to achieve 100% of infection. Forty hours later, cells were prepared, and mitochondria were purified with the Human Mitochondria Isolation Kit (Miltenyi Biotec) according to the manufacturer’s instructions. Briefly, were washed twice with cold PBS and counted. Ten million cells were then lysed with a Dounce homogenizer on ice in the kit lysis buffer supplemented with EDTA-free protease inhibitors (Roche). Cell homogenates were subjected to immunoprecipitation using magnetic beads-coupled anti-TOMM22 antibodies. Isolated mitochondria were centrifuged at 13,000xg for 2 minutes at 4 °C and resuspended in in 40 µl U/T buffer (6 M urea, 2 M thiourea, 10 mM Hepes (pH 8.0)), and reduction and alkylation carried out with 10 mM DTT and 55 mM iodoacetamide in 50 mM ABC buffer (50 mM NH4HCO3 in water pH 8.0), respectively. For the determination of the whole proteome, 5 x 10^6^ washed cells were lysed in a buffer containing 6 M guanidium chloride and 10 mM Tris(2-carboxyethyl)phosphine (TCEP), 0.1M Tris/HCl (pH 8). Fifty micrograms of cleared protein lysates were reduced/alkylated and peptides were purified on stage tips as described above After digestion with 1 µg LysC (WAKO Chemicals USA) at room temperature for 3 h, the suspension was diluted in ABC buffer, and the protein solution was digested with trypsin (Promega) overnight at room temperature. Peptides were purified on self-assembled stage tips with three C18 Empore filter discs (3M) and analyzed by liquid chromatography coupled to mass spectrometry on a QExactive HF instrument (Thermo Fisher Scientific) as previously described (Scaturro *et al*, 2018).

Raw mass-spectrometry data were processed with MaxQuant software versions 1.5.6.2 using the built-in Andromeda search engine to search against the human proteome (*Homo sapiens*; UniprotKB #UP0000005684; release 2012_02) containing forward and reverse sequences concatenated with the DENV (UniprotKB #P29990) and ZIKV (UniprotKB #KU955593) viral proteins, and the label-free quantitation (LFQ) algorithm as described previously (Holze *et al*, 2018; Tyanova *et al*, 2016a). Additionally, the intensity-based absolute quantification (iBAQ) algorithm and “Match Between Runs” option were used. In MaxQuant, carbamidomethylation was set as fixed and methionine oxidation and N-acetylation as variable modifications, using an initial mass tolerance of 6 ppm for the precursor ion and 0.5 Da for the fragment ions. Search results were filtered with a false discovery rate (FDR) of 0.01 for peptide and protein identifications.

Perseus software version 1.6.10.43 was used to further process the affinity-purification and global proteome dataset. Protein tables were filtered to eliminate the identifications from the reverse database and common contaminants. In analyzing mass spectrometry data, only proteins identified on the basis of at least one peptide and a minimum of 3 quantitation events in at least one experimental group were considered. Significant interactors were determined by Welch’s paired T-tests with permutation-based false discovery rate statistics on LFQ intensities (Global proteomes) or the relative abundance of Mitochondria-enriched proteins after normalization against the corresponding cellular lysates (Mitoproteome) (n=4, (|Log2(fold-change))| ≥ 1.58), −Log10(P-value) ≥ 2.). We performed 250 permutations, and the FDR threshold was set at 0.05. The parameter S0 was set at 1 to separate background from specifically enriched interactors (mitoproteome), or to define significantly up-or down-regulated proteins in pairwise comparisons (global proteome). (Tyanova *et al*, 2016b) UniprotKB accession codes of protein groups and proteins associated with Krebs cycle and electron transport chain identified by mass spectrometry, and their respective LFQ intensities, normalized ratios and significance values are provided in Suppl. Table 1. The complete list of identified proteins will be published alongside a different study.

### GC-MS metabolomic analyses

For infection experiments, Huh7.5 cells were infected with DENV2 16681s (MOI=2), ZIKV H/PF/2013 (MOI=10) or ZIKV MR766 (MOI=2) to achieve 100% of infection, and collected two days post-infection for metabolite preparation. For 2K-NS4B overexpression experiments, Huh7.5 cells were transduced with NS4B-expressing lentiviruses (MOI=2). The day after, the medium was replaced and puromycin was added to a final concentration of 1 µg/mL. Three days later (*i.e.*, 4 days post-transduction), selected transduced cells were washed on ice three times with a cold and filtered isotonic solution (0.9% NaCl). Cells were then quickly collected with 800 µL of 80% MS-grade methanol which was stored at −80 °C. Samples were stored at −80 °C until metabolite extraction. In parallel, additional replicate samples were generated for cell counting after trypsinization, and quality controls using western blotting to ensure that the infection and NS4B overexpression were successful. Experiments were design so that between 0.75 x 10^6^ and 2 x 10^6^ cells were used for subsequent processing for GC-MS.

Membranes disruption was carried by sonication at 4 °C (2x10 min, 30 sec on, 30 sec off, high setting, Diagenode Bioruptor). Extracts were cleared by centrifugation (15,000 rpm, 10 min, 4 °C) and supernatants were transferred into new tubes containing 1μl 800 ng/μl myristic acid-D27 (Sigma; dissolved in pyridine). Next, they were dried in a cold trap (Labconco) overnight at − 4 °C. Pellets were solubilized in 30 μl pyridine containing methoxyamine-HCl (10 mg/mL, Sigma) by sonication and vortex, and were incubated at RT for 20 min (methoximation). Samples were centrifuged (15,000 rpm, 10 min, RT) and the supernatants were transferred into glass vials containing MTBSTFA (70 μl, Sigma) for derivatization at 70 °C for 1 h. One μL was injected per sample for GC–MS analysis. GC–MS instrumentation and software were all from Agilent. GC–MS methods and analyses are as previously described (Hulea *et al*, 2018). Data analyses were performed using the Chemstation and MassHunter software (Agilent, Santa Clara, USA). Three replicates per experiment for each condition were processed. The included data show the mean and SEM obtained from the analysis of 4 and 3 ©ndependent experiments for virus infection and 2K-4B expression studies, respectively.

### Caspase-Glo 3/7 assay

300,000 Huh7.5 cells were seeded in 6-well plates and transduced with shRNA-expressing lentivirus at a MOI of 4 with 8µg/mL polybrene. Two days post transduction, cells were infected with ZIKV H/PF/2013 (MOI of 20) or left uninfected. Two days post infection, cells were scraped in culture medium, collected, and centrifuged for 1 min at 10,000 rpm. Cell pellets were resuspended in 70 μL of a 50/50% mixture containing PBS and the Caspase-Glo 3/7 reagent (Promega). Lysates were incubated at least 2 hours protected from the light at room temperature. Luminescence was measured in duplicates in white 96-well plates (30 µl/well) with a Spark multi-mode microplate reader (Tecan). All values were background-subtracted and normalized to the shNT-transduced uninfected condition.

### Flow cytometry

300,000 Huh7.5 cells were prepared exactly as in Caspase-Glo 3/7 assays Two days post-infection, cells were detached by trypsin treatment and stained with 25 nM MitoTracker® Orange CM-H_2_ TMRos (Thermo-Fisher) for 30 min at 37 °C followed by a treatment with 4µM CellEvent caspase 3/7 green (Thermo-Fisher) and the amine reactive viability dye LIVE/DEAD aqua fixable stain (Thermo-Fisher) for 30min in the dark at room temperature. Cells were fixed with 2% formaldehyde and permeabilized with 0.1% Triton X-100. To identify ZIKV-infected cells, total cells were stained for ZIKV using a rat polyclonal anti-NS3 antibodies and subsequently with goat anti-rat cross-adsorbed AlexaFluor 647-conjugated secondary antibodies. Cells were stored at 4°C in the dark until flow cytometry processing (performed within 24 hr) and data acquisition with a BD LSRFortessa instrument at the Flow Cytometry Core Facility of INRS. Data analysis was performed using FlowJo version 10.0 software. After setting of singlets, infected Huh7.5 were defined as NS3+ cells and analyzed for active caspase 3/7 expression and LIVE/DEAD signal.

### Statistical analyses

Statistical significance was evaluated by multiple *t*-test using GraphPad Prism 8.0 software. p values < 0.05 were considered significant: ****: p < 0.0001; ***: p < 0.001; **: p < 0.01; * p < 0.05; ns: not significant.

## ACKNOWLEDGEMENTS

We are grateful to Jessy Tremblay at the Confocal Microscopy and Flow Cytometry Facility of INRS-Centre Armand-Frappier for excellent technical assistance during imaging and data acquisition, and Jeannie Mui and Kelly Sears at the McGill University Facility for Electron Microscopy Research for sample preparation and precious assistance with imaging. We thank the GCRC Metabolomics Core Facility (led by Dr. Daina Avizonis), which is supported by the Canada Foundation for Innovation, the Dr. John R. and Clara M. Fraser Memorial Trust, the Terry Fox Foundation (TFF Oncometabolism Team Grant 1048 in partnership with the Fondation du Cancer du Sein du Quebec), and McGill University. We thank Dr. Ralf Bartenschlager (University of Heidelberg) for providing the DENV reverse genetics system, and Dr. Frédérick Antoine Mallette (University of Montréal), Dr. Tom Hobman (University of Alberta), Dr. Patrick Labonté (Institut National de la Recherche Scientifique) and Dr. Anil Kumar (University of Saskatchewan) for generously providing cell lines. We are grateful to the European Virus Archive Global (EVAg) and Dr. Xavier de Lamballerie (Emergence des Pathologies Virales, Aix-Marseille University, France) for providing ZIKV original stocks.

## CONFLICT OF INTEREST

The authors declare that they have no conflicts of interest.

## FUNDING STATEMENT

W.F received PhD fellowships from the Armand-Frappier Foundation and Fonds de la Recherche du Québec-Santé (FRQS). A.A was a recipient of a master’s training fellowship from FRQS. C.M and A.A.S received PhD fellowships from the Armand-Frappier Foundation and the Center of Excellence in Research on Orphan Diseases-Courtois Foundation (CERMO-FC). A.A.S is receiving a PhD fellowship from the Fonds de la Recherche du Québec-Nature et Technologies (FRQNT). L.C.C has received a research scholar (Junior 2) salary support from FRQS. L.H. is the recipient of a research scholar salary support from FRQS (Junior 1) and the L.H. laboratory is supported by a project grant from the Canadian Institutes of Health Research (CIHR; PJT165901) and an operating grant from the Cancer Research Society (CRS; #25350). Work in A.P’s laboratory was funded by an ERC Consolidator grant (ERC-CoG ProDAP, 817798), the German research foundation (PI 1084/4, PI 1084/5 and TRR179/TP10 and TRR237/A07) and KA1-Co-02 “COVIPA” (Helmholtz Association’s Initiative and Networking Fund). Work in P.S laboratory was founded by the Free and Hanseatic City of Hamburg, the German research foundation (SC314/2-1) and the German Federal Ministry of Education and Research (VirMScan). This research was supported by a project grant from the CIHR (PJT153020), awarded to L.C.C.

## AUTHOR CONTRIBUTIONS

W.F : Execution of the vast majority of the experiments, data analysis, manuscript writing; A.A: PLA and TEM, sample preparation for GC-MS; A.A.S: Apoptosis assay optimisation; C.M: RRBP1 primary characterization; N.T and V.A.B.T: Mitochondria purification; C.G, X.L, I.G.R.G and A.L: FACS data acquisition and analysis; P.S and A.P: mass spectrometry and data analysis. Z.N and L.H: Metabolite extraction, GC-MS and data analyses. L.C.C: Study conceptualization, design and funding, data analysis, manuscript editing. All authors have read, reviewed and agreed to this version of the manuscript.

## SUPPLEMENTAL MATERIALS

**Supplemental figure 1:**
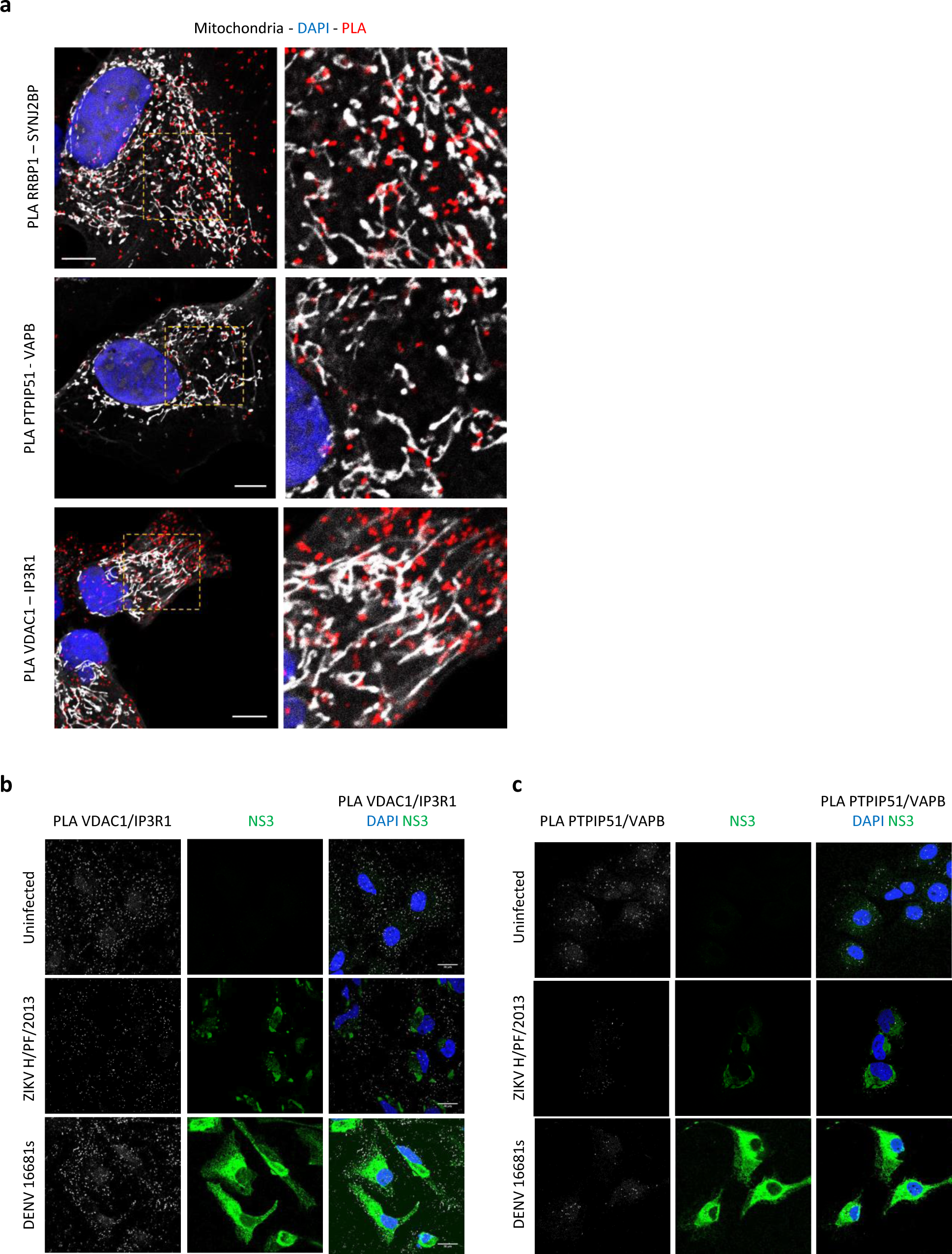
DENV and ZIKV infection decrease the abundance of ERMC tethering complexes. (a) Fixed Huh7.5 cells stably expressingmito-mTurquoise2 (exhibiting fluorescent mitochondria) were subjected to PLAs detecting VDAC1-IP3R1, SYNJ2BP-RRBP1 and VAPB-PTPIP51 interactions. Scale bar = 10 µm. (b,c) Huh7.5 cells were infected with DENV 16681s (MOI=1) or ZIKV H/PF/2013 (MOI=10), or left uninfected. Two days later, cells were fixed and subjected to proximity ligation assays (PLA) to detect (b) VDAC1/IP3R1 or (c) PTPIP51/VAB interactions, and immunostained for NS3 viral protein to identify infected cells. Cells were imaged using confocal microscopy. Representative images are shown.

**Supplemental figure 2:**
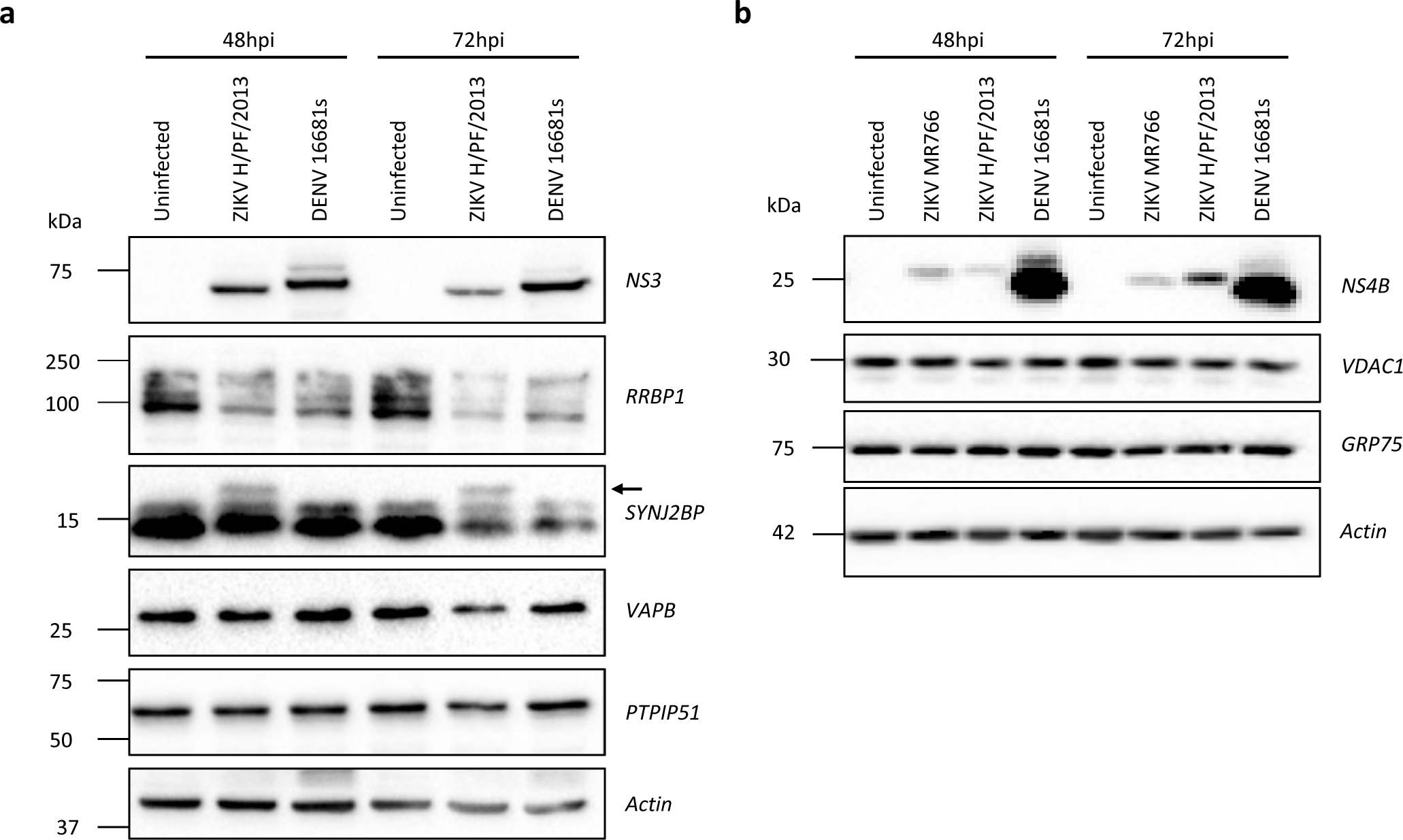
DENV and ZIKV modulates ERMC proteins. (a, b) Huh7.5 cells were infected with DENV 16681s (MOI=1) or ZIKV H/PF/2013 (MOI=10) or left uninfected. Two or three days post-infection, cell extracts were prepared, and the expression levels of the indicated proteins were analyzed by western blotting. The arrow indicates the ZIKV-induced SYNJ2BP protein product.

**Supplemental figure 3:**
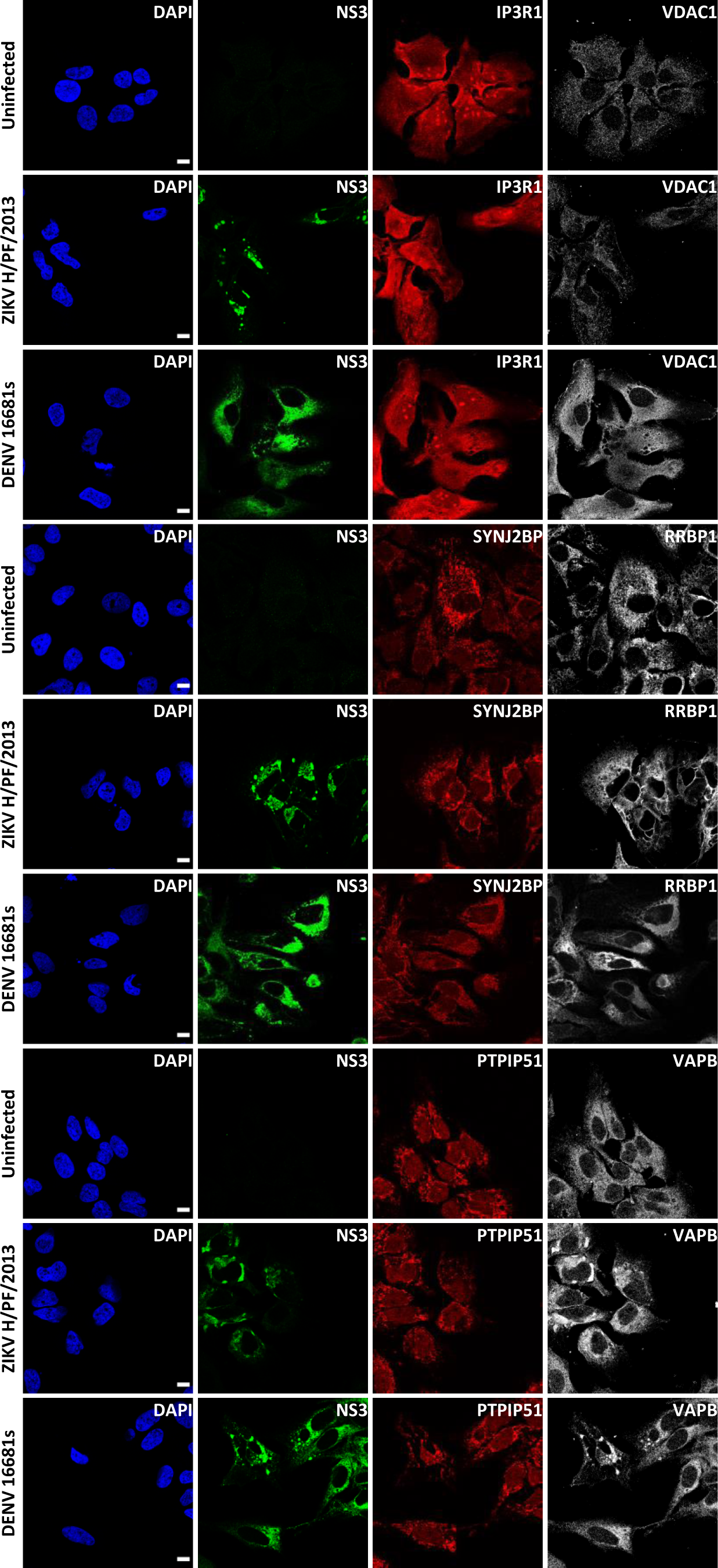
ERMC protein cellular localization in DENV and ZIKV-infected cells. Uninfected and DENV/ZIKV-infected Huh7.5 cells were imaged at two days post-infection by confocal microscopy using antibodies detecting the indicated ERMC proteins. Infected cells were detected with anti-NS3 antibodies.

**Supplemental figure 4:**
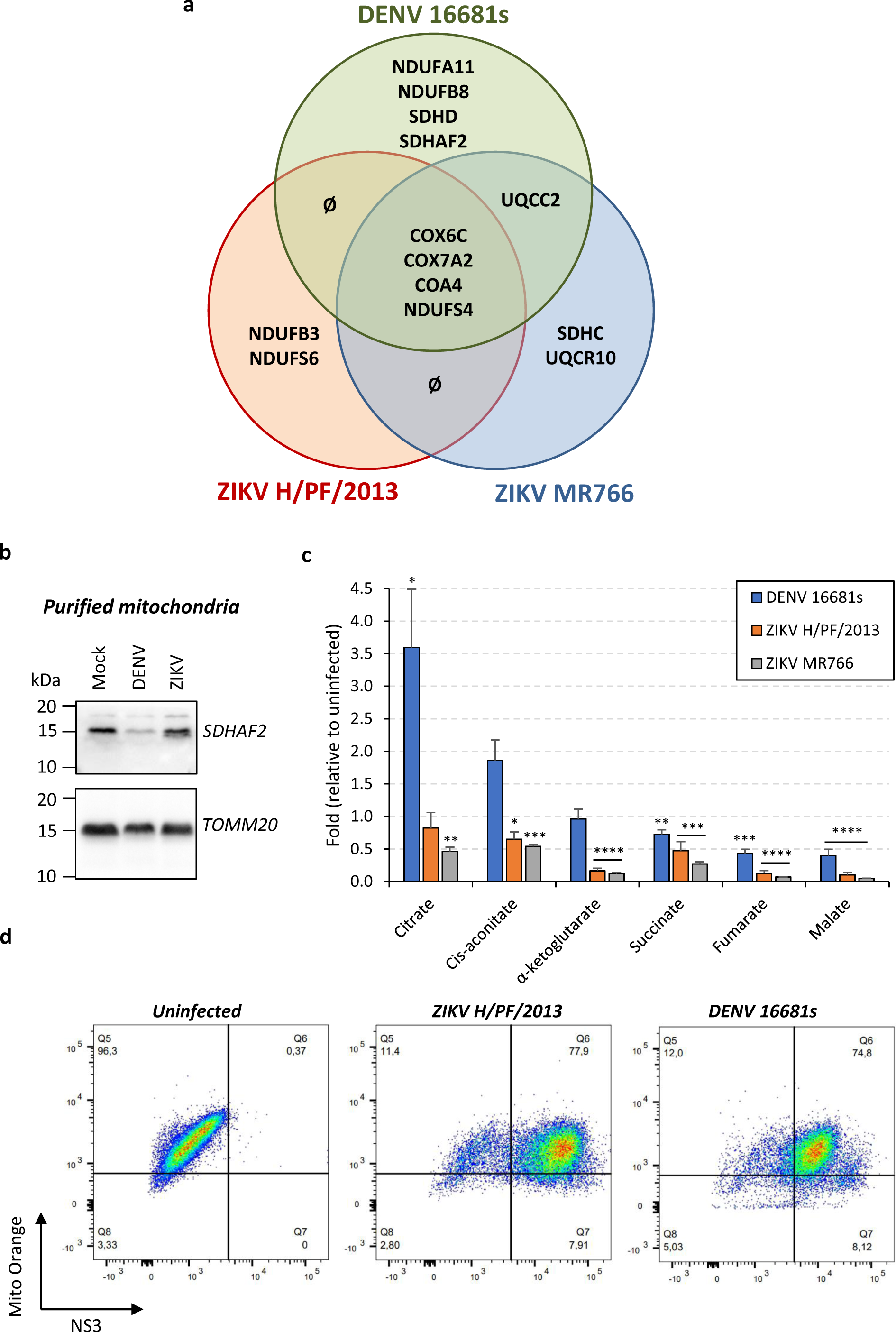
DENV and ZIKV perturb mitochondrial respiratory metabolism. A Venn diagram illustrating the electron transport chain proteins whose mitochondrial abundance is significantly changed when cells are infected with DENV 16681s, ZIKV H/PF/2013 or ZIKV MR766. (b) Huh7.5 cells were infected with DENV 16681s (MOI=2) or ZIKV H/PF/2013 (MOI=10) or left uninfected. Two days later, mitochondria were purified and analyzed by western blotting for their content in SDHAF2 and TOMM20 as control. (c) DENV/ZIKV-infected and control cells were analyzed two days post-infection for the abundance of the Krebs cycle metabolites measured by GC-MS. All values were normalized to the uninfected condition. (d) Cells we treated as in (b) and the mitochondrial potential in uninfected and DENV/ZIKV-infected cells was measured by FACS following staining with MitoTracker Orange CM-H2 TMRos and anti-NS3 antibodies.

**Supplemental figure 5:**
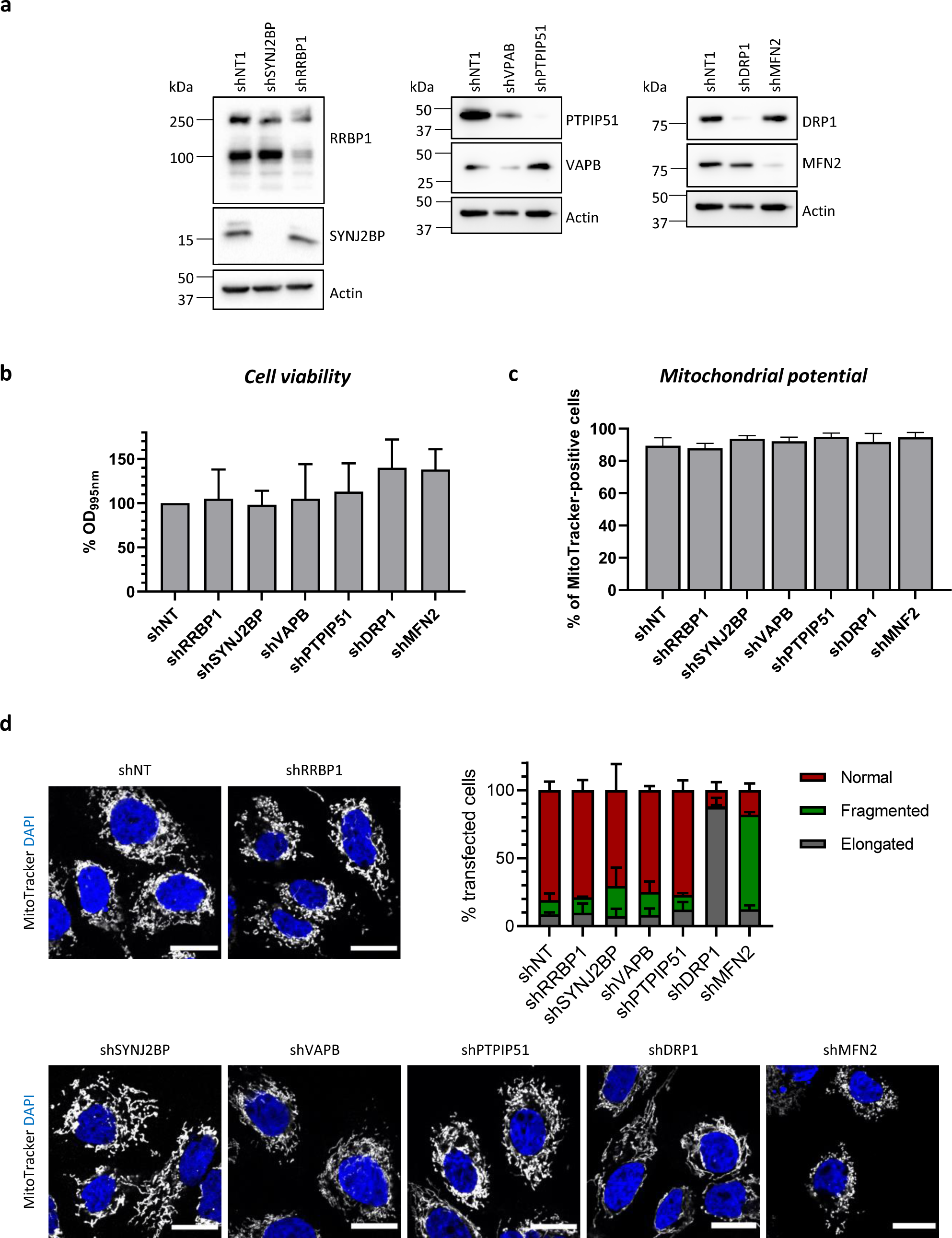
ERMC protein knockdown does not impact mitochondria morphology and integrity. (a-b) Huh7.5 cells were transduced with lentiviruses expressing shRNAs which target the indicated proteins (MOI=5) and selected with puromycin 1 µg/mL. Four days later, cells were either collected for (a) assessing shRNA-mediated knockdown efficiency by western blotting or evaluating the impact of protein depletion on cell viability using MTT assays. (c) Mitochondrial potential upon ERMC protein knockdown was measured by FACS following staining with MitoTracker Orange CM-H2 TMRos. (d) The mitochondrial morphology was analyzed in knocked down cells using confocal microscopy following staining with the MitoTracker Red CMXRos dye. Quantification was made with data from two independent experiments and 75-100 cells per condition per experiment.

**Supplemental figure 6:**
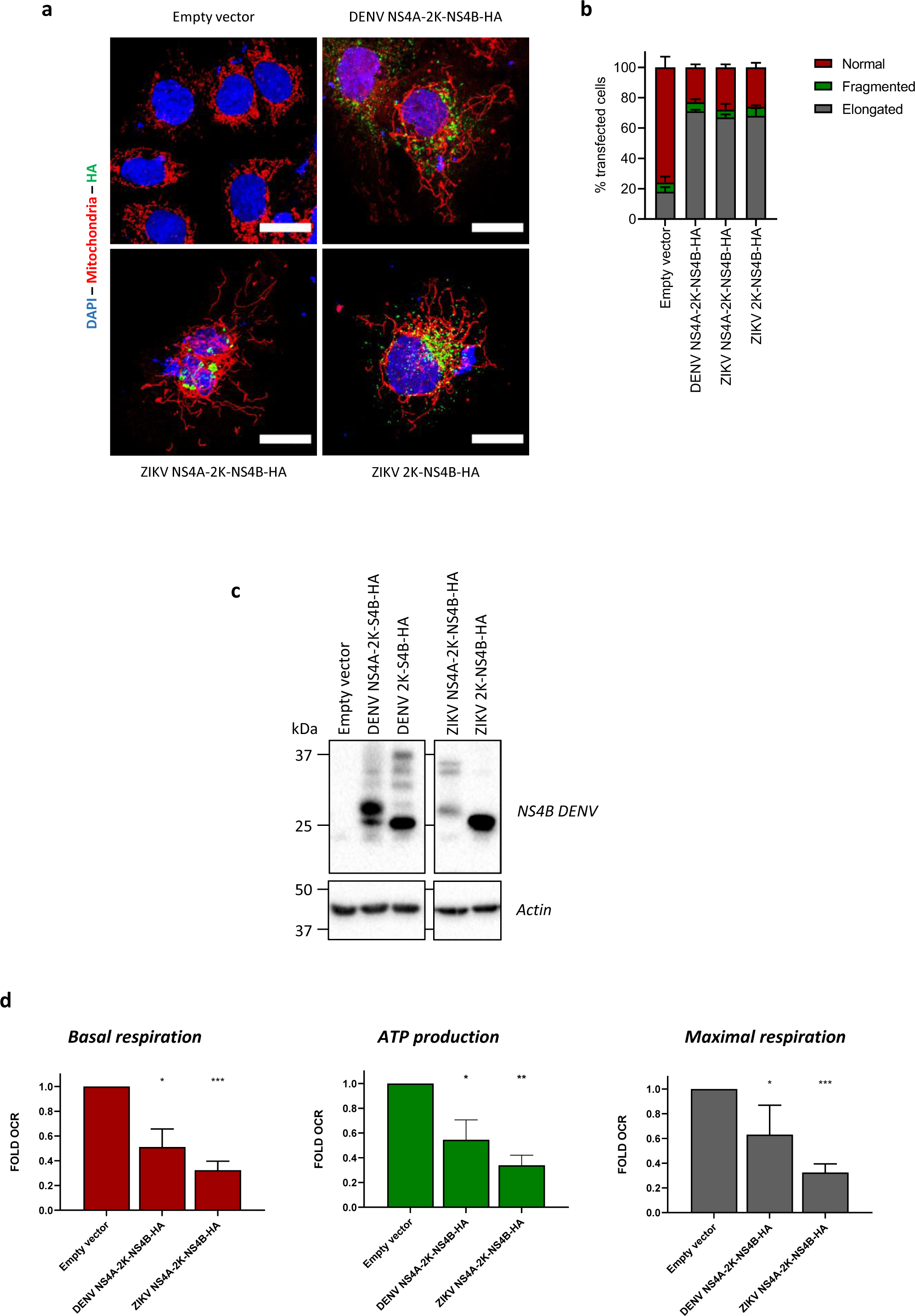
ZIKV NS4B induces mitochondria elongation and DENV/ZIKV NS4B precursors inhibit the mitochondrial respiratory metabolism. (a) Huh7.5/T7 cells expressing the T7 RNA polymerase were transfected with plasmids expressing DENV or ZIKV NS4B constructs. Sixteen hours post-transfection, the mitochondrial morphology was analyzed using confocal microscopy following staining with the MitoTracker Red CMXRos dye and anti-HA antibodies. (b) Quantification of elongated, normal, and fragmented mitochondria in transfected cells from two independent experiments. (c) Huh7.5 cells were transduced with lentiviruses expressing DENV and ZIKV NSA-2K-NS4B which were HA-tagged (MOI=4) and submitted to puromycin selection. Three days post-transduction, cells were analyzed for NS4B expression by western blotting. (d) Huh7.5 cells were transduced with lentiviruses and selected as in (c). Three days post-transduction, equal amounts of transduced living cells were analyzed for their basal respiration, ATP production and maximal respiration using the Seahorse technology. All values were normalised to the control empty vector condition.

## Supplemental table 1

**Targeted proteomic and mitoproteomic analysis of proteins involved in the Krebs cycle and the electron transport chain.** Huh7.5 cells were infected with DENV 16681s (MOI=1) or ZIKV H/PF/2013 (MOI=10), or left uninfected. 2 days later, cell extracts and mitochondria were purified for subsequent quantitative proteomic analysis using mass spectrometry as explained in the Material and Methods section. All values were normalized to the uninfected condition and to total expression levels in case of the mitoproteome. From four independent experiments, the table shows the total relative levels and the mitochondrial enrichment (in Log2) of detected proteins involved in the electron transport chain (ETC) and the Krebs cycle are indicated along with corresponding the p-value (in Log10). A given protein was considered as a protein significantly modulated by ZIKV or DENV in mitoproteome of the cell proteome (depicted with the + symbol) when it met the following hit selection criteria: − Log10(P-value) ≥ 2 and Log2(diff) ≥ 1.58 or ≤ −1.58 (above 3-fold enrichment or depletion, respectively). Hits are indicated in grey (MITO hits or PROT hits columns). Protein LFQ intensity values for each biological replicate are also included in the table. Results are schematically summarized in Figure 3f.

## REFERENCES

1. Anastasia I, Ilacqua N, Raimondi A, Lemieux P, Ghandehari-Alavijeh R, Faure G, Mekhedov SL, Williams KJ, Caicci F, Valle G et al (2021) Mitochondria-rough-ER contacts in the liver regulate systemic lipid homeostasis. Cell Rep 34: 108873

2. Anton A, Mazeaud C, Freppel W, Gilbert C, Tremblay N, Sow AA, Roy M, Rodrigue-Gervais IG, Chatel-Chaix L (2021) Valosin-containing protein ATPase activity regulates the morphogenesis of Zika virus replication organelles and virus-induced cell death. Cell Microbiol: e13302

3. Barbier V, Lang D, Valois S, Rothman AL, Medin CL (2017) Dengue virus induces mitochondrial elongation through impairment of Drp1-triggered mitochondrial fission. Virology 500: 149–160

4. Bhatt S, Gething PW, Brady OJ, Messina JP, Farlow AW, Moyes CL, Drake JM, Brownstein JS, Hoen AG, Sankoh O et al (2013) The global distribution and burden of dengue. Nature 496: 504–507

5. Carod-Artal FJ (2018) Neurological complications of Zika virus infection. Expert Rev Anti Infect Ther 16: 399–410

6. Carpio MA, Means RE, Brill AL, Sainz A, Ehrlich BE, Katz SG (2021) BOK controls apoptosis by Ca(2+) transfer through ER-mitochondrial contact sites. Cell Rep 34: 108827

7. Chatel-Chaix L, Bartenschlager R (2014) Dengue virus-and hepatitis C virus-induced replication and assembly compartments: the enemy inside--caught in the web. Journal of virology 88: 5907–5911

8. Chatel-Chaix L, Cortese M, Romero-Brey I, Bender S, Neufeldt CJ, Fischl W, Scaturro P, Schieber N, Schwab Y, Fischer B et al (2016) Dengue Virus Perturbs Mitochondrial Morphodynamics to Dampen Innate Immune Responses. Cell host & microbe 20: 342–356

9. Chatel-Chaix L, Fischl W, Scaturro P, Cortese M, Kallis S, Bartenschlager M, Fischer B, Bartenschlager R (2015) A Combined Genetic-Proteomic Approach Identifies Residues within Dengue Virus NS4B Critical for Interaction with NS3 and Viral Replication. Journal of virology 89: 7170–7186

10. Chen Q, Gouilly J, Ferrat YJ, Espino A, Glaziou Q, Cartron G, El Costa H, Al-Daccak R, Jabrane-Ferrat N (2020) Metabolic reprogramming by Zika virus provokes inflammation in human placenta. Nature communications 11: 2967

11. Cohen S, Valm AM, Lippincott-Schwartz J (2018) Interacting organelles. Curr Opin Cell Biol 53: 84–91

12. Colombini M (2016) The VDAC channel: Molecular basis for selectivity. Biochimica et biophysica acta 1863: 2498–2502

13. Cook KC, Tsopurashvili E, Needham JM, Thompson SR, Cristea IM (2022) Restructured membrane contacts rewire organelles for human cytomegalovirus infection. Nature communications 13: 4720

14. Cortese M, Goellner S, Acosta EG, Neufeldt CJ, Oleksiuk O, Lampe M, Haselmann U, Funaya C, Schieber N, Ronchi P et al (2017) Ultrastructural Characterization of Zika Virus Replication Factories. Cell Rep 18: 2113–2123

15. Csordas G, Renken C, Varnai P, Walter L, Weaver D, Buttle KF, Balla T, Mannella CA, Hajnoczky G (2006) Structural and functional features and significance of the physical linkage between ER and mitochondria. J Cell Biol 174: 915–921

16. De Vos KJ, Morotz GM, Stoica R, Tudor EL, Lau KF, Ackerley S, Warley A, Shaw CE, Miller CC (2012) VAPB interacts with the mitochondrial protein PTPIP51 to regulate calcium homeostasis. Hum Mol Genet 21: 1299–1311

17. Deng L, Adachi T, Kitayama K, Bungyoku Y, Kitazawa S, Ishido S, Shoji I, Hotta H (2008) Hepatitis C virus infection induces apoptosis through a Bax-triggered, mitochondrion-mediated, caspase 3-dependent pathway. Journal of virology 82: 10375–10385

18. Duan X, Li S, Holmes JA, Tu Z, Li Y, Cai D, Liu X, Li W, Yang C, Jiao B et al (2018) MicroRNA 130a Regulates both Hepatitis C Virus and Hepatitis B Virus Replication through a Central Metabolic Pathway. Journal of virology 92

19. Duan Y, Wang X, Sun K, Lin Y, Wang X, Chen K, Yang G, Wang X, Du C (2022) SYNJ2BP Improves the Production of Lentiviral Envelope Protein by Facilitating the Formation of Mitochondrion-Associated Endoplasmic Reticulum Membrane. Journal of virology 96: e0054922

20. El-Bacha T, Midlej V, Pereira da Silva AP, Silva da Costa L, Benchimol M, Galina A, Da Poian AT (2007) Mitochondrial and bioenergetic dysfunction in human hepatic cells infected with dengue 2 virus. Biochimica et biophysica acta 1772: 1158–1166

21. Elmore S (2007) Apoptosis: a review of programmed cell death. Toxicol Pathol 35: 495–516

22. Fischl W, Bartenschlager R (2013) High-throughput screening using dengue virus reporter genomes. Methods Mol Biol 1030: 205–219

23. Freppel W, Mazeaud C, Chatel-Chaix L (2018) Production, Titration and Imaging of Zika Virus in Mammalian Cells. Bio-Protocol 8

24. Friedman JR, Lackner LL, West M, DiBenedetto JR, Nunnari J, Voeltz GK (2011) ER tubules mark sites of mitochondrial division. Science 334: 358–362

25. Fujimoto M, Hayashi T (2011) New insights into the role of mitochondria-associated endoplasmic reticulum membrane. Int Rev Cell Mol Biol 292: 73–117

26. Gack MU, Diamond MS (2016) Innate immune escape by Dengue and West Nile viruses. Curr Opin Virol 20: 119–128

27. Gack MU, Shin YC, Joo CH, Urano T, Liang C, Sun L, Takeuchi O, Akira S, Chen Z, Inoue S et al (2007) TRIM25 RING-finger E3 ubiquitin ligase is essential for RIG-I-mediated antiviral activity. Nature 446: 916–920

28. Giamogante F, Poggio E, Barazzuol L, Covallero A, Cali T (2021) Apoptotic signals at the endoplasmic reticulum-mitochondria interface. Adv Protein Chem Struct Biol 126: 307–343

29. Gillespie LK, Hoenen A, Morgan G, Mackenzie JM (2010) The endoplasmic reticulum provides the membrane platform for biogenesis of the flavivirus replication complex. Journal of virology 84: 10438–10447

30. Gomez-Suaga P, Paillusson S, Stoica R, Noble W, Hanger DP, Miller CCJ (2017) The ER-Mitochondria Tethering Complex VAPB-PTPIP51 Regulates Autophagy. Current biology : CB 27: 371–385

31. Grant D, Tan GK, Qing M, Ng JK, Yip A, Zou G, Xie X, Yuan Z, Schreiber MJ, Schul W et al (2011) A single amino acid in nonstructural protein NS4B confers virulence to dengue virus in AG129 mice through enhancement of viral RNA synthesis. Journal of virology 85: 7775–7787

32. Hamasaki M, Furuta N, Matsuda A, Nezu A, Yamamoto A, Fujita N, Oomori H, Noda T, Haraguchi T, Hiraoka Y et al (2013) Autophagosomes form at ER-mitochondria contact sites. Nature 495: 389–393

33. Hartmann J, Verkhratsky A (1998) Relations between intracellular Ca2+ stores and store-operated Ca2+ entry in primary cultured human glioblastoma cells. J Physiol 513 (Pt 2): 411–424

34. Holze C, Michaudel C, Mackowiak C, Haas DA, Benda C, Hubel P, Pennemann FL, Schnepf D, Wettmarshausen J, Braun M et al (2018) Oxeiptosis, a ROS-induced caspase-independent apoptosis-like cell-death pathway. Nat Immunol 19: 130–140

35. Horner SM, Liu HM, Park HS, Briley J, Gale M, Jr. (2011) Mitochondrial-associated endoplasmic reticulum membranes (MAM) form innate immune synapses and are targeted by hepatitis C virus. Proceedings of the National Academy of Sciences of the United States of America 108: 14590–14595

36. Horner SM, Wilkins C, Badil S, Iskarpatyoti J, Gale M, Jr. (2015) Proteomic analysis of mitochondrial-associated ER membranes (MAM) during RNA virus infection reveals dynamic changes in protein and organelle trafficking. PloS one 10: e0117963

37. Hulea L, Gravel SP, Morita M, Cargnello M, Uchenunu O, Im YK, Lehuede C, Ma EH, Leibovitch M, McLaughlan S et al (2018) Translational and HIF-1alpha-Dependent Metabolic Reprogramming Underpin Metabolic Plasticity and Responses to Kinase Inhibitors and Biguanides. Cell Metab 28: 817–832 e818

38. Hung V, Lam SS, Udeshi ND, Svinkina T, Guzman G, Mootha VK, Carr SA, Ting AY (2019) Correction: Proteomic mapping of cytosol-facing outer mitochondrial and ER membranes in living human cells by proximity biotinylation. Elife 8

39. Jenner A, Pena-Blanco A, Salvador-Gallego R, Ugarte-Uribe B, Zollo C, Ganief T, Bierlmeier J, Mund M, Lee JE, Ries J et al (2022) DRP1 interacts directly with BAX to induce its activation and apoptosis. EMBO J 41: e108587

40. Junjhon J, Pennington JG, Edwards TJ, Perera R, Lanman J, Kuhn RJ (2014) Ultrastructural characterization and three-dimensional architecture of replication sites in dengue virus-infected mosquito cells. Journal of virology 88: 4687–4697

41. Kaptein SJF, Goethals O, Kiemel D, Marchand A, Kesteleyn B, Bonfanti JF, Bardiot D, Stoops B, Jonckers THM, Dallmeier K et al (2021) Publisher Correction: A pan-serotype dengue virus inhibitor targeting the NS3-NS4B interaction. Nature 599: E2

42. Kelly EP, Puri B, Sun W, Falgout B (2010) Identification of mutations in a candidate dengue 4 vaccine strain 341750 PDK20 and construction of a full-length cDNA clone of the PDK20 vaccine candidate. Vaccine 28: 3030–3037

43. Ledur PF, Karmirian K, Pedrosa C, Souza LRQ, Assis-de-Lemos G, Martins TM, Ferreira J, de Azevedo Reis GF, Silva ES, Silva D et al (2020) Zika virus infection leads to mitochondrial failure, oxidative stress and DNA damage in human iPSC-derived astrocytes. Scientific reports 10: 1218

44. Liu-Helmersson J, Quam M, Wilder-Smith A, Stenlund H, Ebi K, Massad E, Rocklov J (2016) Climate Change and Aedes Vectors: 21st Century Projections for Dengue Transmission in Europe. EBioMedicine 7: 267##x2013;277

45. Liu HM, Loo YM, Horner SM, Zornetzer GA, Katze MG, Gale M, Jr. (2012) The mitochondrial targeting chaperone 14-3-3epsilon regulates a RIG-I translocon that mediates membrane association and innate antiviral immunity. Cell host & microbe 11: 528–537

46. Loo YM, Gale M, Jr. (2011) Immune signaling by RIG-I-like receptors. Immunity 34: 680–692

47. Manokaran G, Finol E, Wang C, Gunaratne J, Bahl J, Ong EZ, Tan HC, Sessions OM, Ward AM, Gubler DJ et al (2015) Dengue subgenomic RNA binds TRIM25 to inhibit interferon expression for epidemiological fitness. Science 350: 217–221

48. Mazeaud C, Anton A, Pahmeier F, Sow AA, Cerikan B, Freppel W, Cortese M, Bartenschlager R, Chatel-Chaix L (2021) The Biogenesis of Dengue Virus Replication Organelles Requires the ATPase Activity of Valosin-Containing Protein. Viruses 13

49. Mazeaud C, Freppel W, Chatel-Chaix L (2018) The Multiples Fates of the Flavivirus RNA Genome During Pathogenesis. Frontiers in genetics 9: 595

50. Miller S, Kastner S, Krijnse-Locker J, Buhler S, Bartenschlager R (2007) The non-structural protein 4A of dengue virus is an integral membrane protein inducing membrane alterations in a 2K-regulated manner. The Journal of biological chemistry 282: 8873–8882

51. Miorin L, Romero-Brey I, Maiuri P, Hoppe S, Krijnse-Locker J, Bartenschlager R, Marcello A (2013) Three-dimensional architecture of tick-borne encephalitis virus replication sites and trafficking of the replicated RNA. Journal of virology 87: 6469–6481

52. Montessuit S, Somasekharan SP, Terrones O, Lucken-Ardjomande S, Herzig S, Schwarzenbacher R, Manstein DJ, Bossy-Wetzel E, Basanez G, Meda P et al (2010) Membrane remodeling induced by the dynamin-related protein Drp1 stimulates Bax oligomerization. Cell 142: 889–901

53. Moore CA, Staples JE, Dobyns WB, Pessoa A, Ventura CV, Fonseca EB, Ribeiro EM, Ventura LO, Neto NN, Arena JF et al (2017) Characterizing the Pattern of Anomalies in Congenital Zika Syndrome for Pediatric Clinicians. JAMA Pediatr 171: 288–295

54. Moquin SA, Simon O, Karuna R, Lakshminarayana SB, Yokokawa F, Wang F, Saravanan C, Zhang J, Day CW, Chan K et al (2021) NITD-688, a pan-serotype inhibitor of the dengue virus NS4B protein, shows favorable pharmacokinetics and efficacy in preclinical animal models. Sci Transl Med 13

55. Oettinghaus B, D’Alonzo D, Barbieri E, Restelli LM, Savoia C, Licci M, Tolnay M, Frank S, Scorrano L (2016) DRP1-dependent apoptotic mitochondrial fission occurs independently of BAX, BAK and APAF1 to amplify cell death by BID and oxidative stress. Biochimica et biophysica acta 1857: 1267–1276

56. Orrenius S, Zhivotovsky B, Nicotera P (2003) Regulation of cell death: the calcium-apoptosis link. Nature reviews Molecular cell biology 4: 552–565

57. Park JH, Ko J, Hwang J, Koh HC (2015) Dynamin-related protein 1 mediates mitochondria-dependent apoptosis in chlorpyrifos-treated SH-SY5Y cells. Neurotoxicology 51: 145–157

58. Paul D, Bartenschlager R (2015) Flaviviridae Replication Organelles: Oh, What a Tangled Web We Weave. Annu Rev Virol 2: 289–310

59. Pourcelot M, Arnoult D (2014) Mitochondrial dynamics and the innate antiviral immune response. FEBS J 281: 3791–3802

60. Riedl W, Acharya D, Lee JH, Liu G, Serman T, Chiang C, Chan YK, Diamond MS, Gack MU (2019) Zika Virus NS3 Mimics a Cellular 14-3-3-Binding Motif to Antagonize RIG-I-and MDA5-Mediated Innate Immunity. Cell host & microbe 26: 493–503 e496

61. Ryan SJ, Carlson CJ, Mordecai EA, Johnson LR (2019) Global expansion and redistribution of Aedes-borne virus transmission risk with climate change. PLoS neglected tropical diseases 13: e0007213

62. Scaturro P, Stukalov A, Haas DA, Cortese M, Draganova K, Plaszczyca A, Bartenschlager R, Gotz M, Pichlmair A (2018) An orthogonal proteomic survey uncovers novel Zika virus host factors. Nature 561: 253–257

63. Schwarz DS, Blower MD (2016) The endoplasmic reticulum: structure, function and response to cellular signaling. Cell Mol Life Sci 73: 79–94

64. Serman TM, Gack MU (2019) Evasion of Innate and Intrinsic Antiviral Pathways by the Zika Virus. Viruses 11

65. Shan C, Xie X, Muruato AE, Rossi SL, Roundy CM, Azar SR, Yang Y, Tesh RB, Bourne N, Barrett AD et al (2016) An Infectious cDNA Clone of Zika Virus to Study Viral Virulence, Mosquito Transmission, and Antiviral Inhibitors. Cell host & microbe 19: 891–900

66. Stoica R, De Vos KJ, Paillusson S, Mueller S, Sancho RM, Lau KF, Vizcay-Barrena G, Lin WL, Xu YF, Lewis J et al (2014) ER-mitochondria associations are regulated by the VAPB-PTPIP51 interaction and are disrupted by ALS/FTD-associated TDP-43. Nature communications 5: 3996

67. Suresh SN (2019) Endoplasmic reticulum mitochondria contacts modulate apoptosis of renal cells and its implications in diabetic neuropathy. EBioMedicine 44: 24–25

68. Tremblay N, Freppel W, Sow AA, Chatel-Chaix L (2019) The Interplay between Dengue Virus and the Human Innate Immune System: A Game of Hide and Seek. Vaccines 7

69. Tyanova S, Temu T, Cox J (2016a) The MaxQuant computational platform for mass spectrometry-based shotgun proteomics. Nat Protoc 11: 2301–2319

70. Tyanova S, Temu T, Sinitcyn P, Carlson A, Hein MY, Geiger T, Mann M, Cox J (2016b) The Perseus computational platform for comprehensive analysis of (prote)omics data. Nat Methods 13: 731–740

71. van Cleef KW, Overheul GJ, Thomassen MC, Kaptein SJ, Davidson AD, Jacobs M, Neyts J, van Kuppeveld FJ, van Rij RP (2013) Identification of a new dengue virus inhibitor that targets the viral NS4B protein and restricts genomic RNA replication. Antiviral Res 99: 165–171

72. Vance JE (2015) Phospholipid synthesis and transport in mammalian cells. Traffic 16: 1–18

73. Verfaillie T, Rubio N, Garg AD, Bultynck G, Rizzuto R, Decuypere JP, Piette J, Linehan C, Gupta S, Samali A et al (2012) PERK is required at the ER-mitochondrial contact sites to convey apoptosis after ROS-based ER stress. Cell death and differentiation 19: 1880–1891

74. Welsch S, Miller S, Romero-Brey I, Merz A, Bleck CK, Walther P, Fuller SD, Antony C, Krijnse-Locker J, Bartenschlager R (2009) Composition and three-dimensional architecture of the dengue virus replication and assembly sites. Cell host & microbe 5: 365–375

75. WHO (2022) Fact sheets: Dengue and severe dengue, 10 January 2022. https://www.whoint/news-room/fact-sheets/detail/dengue-and-severe-dengue

76. Xie X, Wang QY, Xu HY, Qing M, Kramer L, Yuan Z, Shi PY (2011) Inhibition of dengue virus by targeting viral NS4B protein. Journal of virology 85: 11183–11195

77. Yang S, Gorshkov K, Lee EM, Xu M, Cheng YS, Sun N, Soheilian F, de Val N, Ming G, Song H et al (2020) Zika Virus-Induced Neuronal Apoptosis via Increased Mitochondrial Fragmentation. Frontiers in microbiology 11: 598203

78. Yeo HK, Park TH, Kim HY, Jang H, Lee J, Hwang GS, Ryu SE, Park SH, Song HK, Ban HS et al (2021) Phospholipid transfer function of PTPIP51 at mitochondria-associated ER membranes. EMBO reports 22: e51323

79. Zou J, Lee le T, Wang QY, Xie X, Lu S, Yau YH, Yuan Z, Geifman Shochat S, Kang C, Lescar J et al (2015) Mapping the Interactions between the NS4B and NS3 proteins of dengue virus. Journal of virology 89: 3471–3483

80. Zou J, Xie X, Lee le T, Chandrasekaran R, Reynaud A, Yap L, Wang QY, Dong H, Kang C, Yuan Z et al (2014) Dimerization of flavivirus NS4B protein. Journal of virology 88: 3379–3391

